# Relationship between vestibular hair cell loss and deficits in two anti-gravity reflexes in the rat

**DOI:** 10.1101/2020.12.21.423804

**Authors:** Alberto F. Maroto, Alejandro Barrallo-Gimeno, Jordi Llorens

## Abstract

The tail-lift reflex and the air-righting reflex in rats are anti-gravity reflexes that depend on vestibular function. To begin identifying their cellular basis, this study examined the relationship between reflex loss and the graded lesions caused in the vestibular sensory epithelia by varying doses of an ototoxic compound. After ototoxic exposure, we recorded these reflexes using high speed video. The movies were used to obtain objective measures of the reflexes: the minimum angle formed by the nose, the back of the neck and the base of the tail during the tail-lift maneuver and the time to right in the air-righting test. The vestibular sensory epithelia were then collected from the rats and used to estimate the loss of type I (HCI), type II (HCII) and all hair cells (HC) in both central and peripheral parts of the crista, utricle, and saccule. As expected, tail-lift angles decreased, and air-righting times increased, while the numbers of HCs remaining in the epithelia decreased in a dose-dependent manner. The results demonstrated greater sensitivity of HCI compared to HCII to the IDPN ototoxicity, as well as a relative resiliency of the saccule compared to the crista and utricle. Comparing the functional measures with the cell counts, we observed that loss of the tail-lift reflex associates better with HCI than with HCII loss. In contrast, most HCI in the crista and utricle were lost before air-righting times increased. These data suggest that these reflexes depend on the function of non-identical populations of vestibular HCs.

## 1. INTRODUCTION

The vestibular system is the sensory system that encodes head accelerations. As generated by gravity and movements of the body or the head alone, these accelerations can be linear or angular and of a wide diversity of magnitudes. The vestibular system allows for a precise encoding of this large variability of stimuli, thanks to the diversity of functional subsystems that it contains. These are based on the existence of diverse end-organs, types of transducing cells, primary afferent terminals, and physiological properties of the primary neurons (Baird et al., 1988; Desai et al., 2005a, 2005b; Eatock and Songer, 2011; Curthoys et al., 2017; Eatock, 2018).

In mammals, each inner ear has five vestibular sensory epithelia, one crista in each of three semicircular canals, and two maculae in the utricle and saccule (Fig 1A). In each of these end-organs, the cellular architecture defines two broad areas, that of the crista central zones and macular striolae, and that of the peripheral zones (Fig 1B). There are two types of sensory transducing cells, known as hair cells (HCs). Type I HCs (HCI) have an amphora-like shape, whereas type II HCs (HCII) are more cylindrical in their upper part (Fig 1C). HCs are presynaptic to terminals of the afferent neurons of the vestibular ganglion. There are two types of terminals: calyx terminals encase HCI up to their neck, whereas HCII are contacted by standard bouton terminals. These two types of terminals define three types of afferents: calyx-only afferents that exclusively form calyx terminals in the central zones of the epithelia, bouton-only afferents, and dimorphic afferents that form both calyx terminals and bouton terminals. Physiologically, afferents show diverse resting spike timing. Afferents with highly regular timing are more abundant in the peripheral zones of the receptors, while afferents with more irregular spike timing concentrate towards the central crista or the striola of the maculae. On stimulation, more regular or irregular afferents show, respectively, more tonic or transient responses. Thus, irregular afferents are calyx-only or dimorphic and regular afferents are dimorphic or bouton-only, so both HCI and HCII provide input to afferents of diverse morphological and physiological properties. This arrangement greatly complicates the analysis of the roles played by the diverse zones and HC types in vestibular physiology (Eatock, 2018).

**Figure 1.**
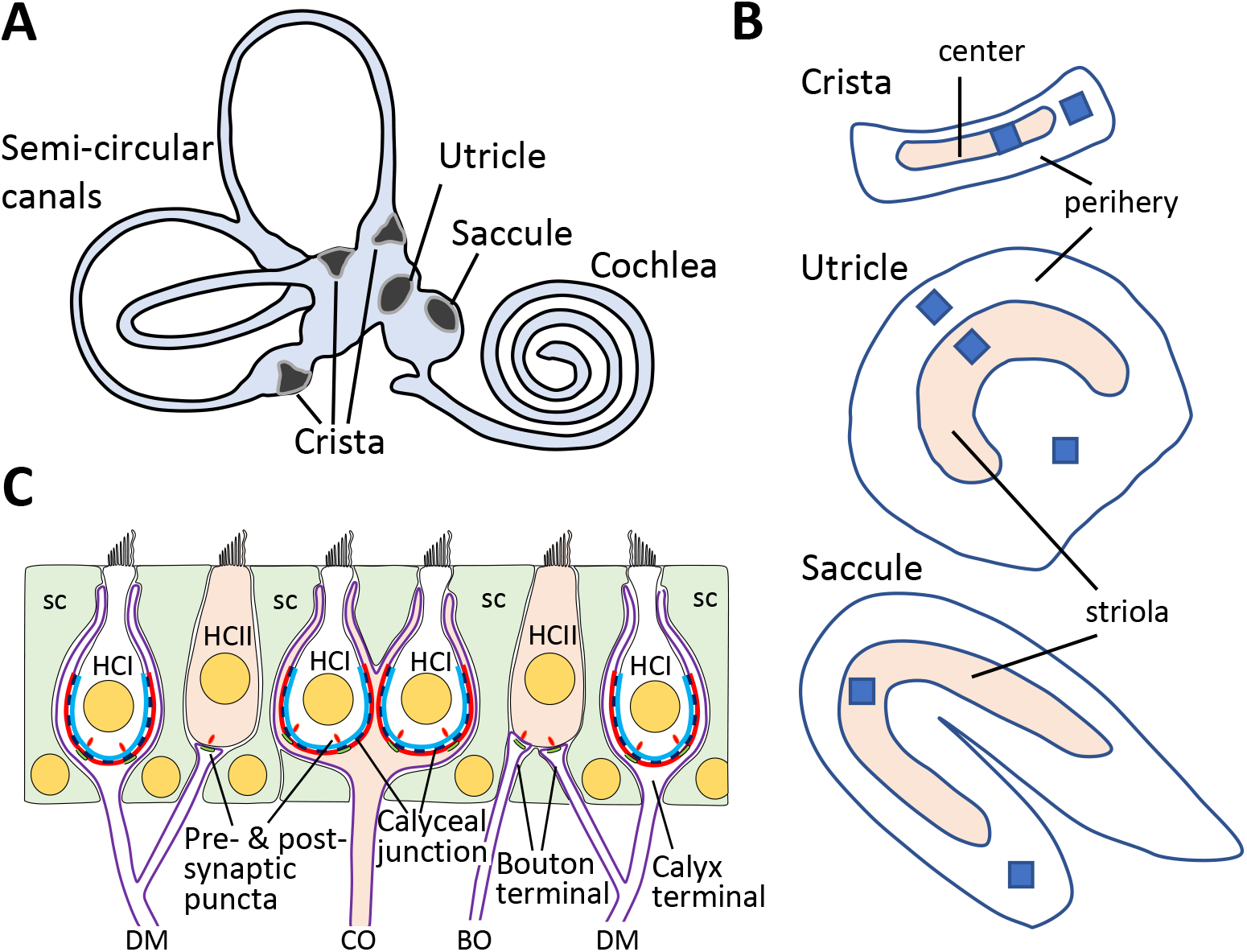
A. Schematic drawing of the mammalian inner ear, showing the five vestibular sensory epithelia, three cristae, the utricle and the saccule. B. In each vestibular epithelia a central zone (named striola for the utricle and saccule) can be differentiated from the peripheral zones by several histological and physiological criteria. Squares show the approximate size and location of the seven areas imaged for cell counting: crista center, crista periphery, utricle lateral periphery (external to the striola), utricle striola, utricle medial periphery (internal to the striola), saccule striola and saccule periphery. C. Cell types, afferent terminals and structural characteristics of the vestibular epithelium. The epithelium consists of type I hair cells (HCI), type II hair cells (HCII) and supporting cells (sc). Calyceal junctions characterize the contacts between the basolateral part of the HCIs and the facing membrane of the calyces encasing them, as represented by the cyan and red lines, respectively. HCII are contacted by bouton terminals. There are three types of afferents: dimorphic (DM), forming both calyx and bouton terminals, calyx-only (CO) and bouton-only (BO) afferents. Calyx-only afferents characterize the central/striola regions. The HCs contain pre-synaptic structures (seen as puncta when immunolabelled) and the afferents contain post-synaptic densities (post-synaptic puncta).

Vestibular information is used for a variety of purposes, including gaze stabilization, motor control and cognitive functions. The understanding of the precise role of each vestibular subsystem to each function is far from being complete. This includes a limited understanding of the substrate underneath vestibular dysfunction symptoms. To tackle these limitations, there is a need for a multiplicity of tests of vestibular function for which the anatomical, cellular, physiological, and molecular substrates are known, including the correspondence between human and animal measures. In clinical settings, testing of vestibular function is increasingly performed through the video head impulse test (vHIT) and assessment of the ocular and cervical vestibular-evoked myogenic potentials (oVEMP and cVEMP, respectively). All of these are measures of reflex responses elicited by fast stimuli. Thus, the vHIT test allows to individually test the six semicircular canals of the patient at high rotational accelerations that impede interference of the optokinetic reflex (Halmagyi et al., 2017). In the case of VEMPs, available evidence indicates that they reflect mainly the function of afferents from the striolar zone of the utricle (oVEMPs) or the saccule (cVEMPS) (Curthoys et al., 2012; Corneil and Camp, 2018; Curthoys et al., 2017, 2018; Ono et al., 2020).

In comparison to clinical tests, rodent tests of vestibular function remain underdeveloped. The better developed test is the evaluation of the vestibulo-ocular reflex (VOR), with most examples involving mice stimulated with low angular velocities (Beraneck et al., 2012; De Jeu and De Zeeuw, 2012; Imai et al., 2016). Another example is the recording of vestibular evoked potential responses to jerk stimuli in mice, recently shown to be defective in mice with compromised formation of the striolar/central zones (Ono et al., 2020). We have recently described the use of high-speed video recording to obtain objective and fully quantitative measures of two anti-gravitational reflexes in the rat, the tail-lift reflex, and the air righting reflex (Martins-Lopes et al., 2019). The first of these tests evaluates the trunk extension reflex shown by rats when lifted by the tail, by measuring the minimum angle formed by the nose, the back of the neck and the base of the tail during the lift maneuver. This reflex is lost in vestibular deficient animals, which show instead ventral curling, and therefore reduced angles (Hunt et al. 1987; Pellis et al., 1991). The second test evaluates the righting-in-the-air response by measuring the time taken by the head of the rat to right after the animal is released in supine position in the air to fall on a foam cushion. The role of vestibular input in this reflex is well documented (Pellis et al., 1989). Our previous study (Martins-Lopes et al., 2019) demonstrated that the minimum tail-lift angle and the air-righting time provide good measures of vestibular function loss following ototoxic insults. In the present study, we study the relationship between reflex loss and loss of sensory HCs in the vestibular epithelia following the ototoxic insult. To gain insight into the contribution of the different HC types and epithelial zones to the reflexes, we separately estimated the loss of HCI, HCII and all HCs in central/striolar zones and peripheral zones of all three vestibular end-organs. The results suggest that these two reflexes depend on non-identical populations of HCs and that the tail-lift reflex is altered at earlier stages of HC loss than the air-righting reflex.

## 2. MATERIAL AND METHODS

### 2.1. Animals and treatments

This study used three lots of male adult (8-9-week-old) Long-Evans rats (n=24, 25 and 24), obtained from Janvier Labs (Le-Genest-Saint-Isle, France). The animals were housed in groups of two or three in standard cages (215 x 465 x 145 mm) with wood shavings as bedding. Housing conditions were a 12:12 h light:dark cycle (07: 30-19:30 h), a temperature of 22 °C ± 2°C, and free access to food pellets (TEKLAD 2014, Harlan Laboratories, Sant Feliu de Codines, Spain). The rats were housed for acclimatization for six days before starting the experiments. During experiments, the animals were regularly weighed and evaluated for overall toxicity to limit suffering according to ethical criteria. The use of the animals was in accordance with EU Directive 2010/63, following Law 5/1995 and Act 214/1997 of the Generalitat de Catalunya, and Law 6/2013 and Act 53/2013 of the Gobierno de España. This included compulsory approval by the Ethics Committee on Animal Experimentation of the Universitat de Barcelona.

HC loss was induced by exposure to 3,3’-iminodipropionitrile (IDPN), a well characterized ototoxic compound (Crofton and Knigth, 1991; Llorens et al., 1993; Llorens and Demêmes, 1994; Crofton et al., 1994; Llorens and Rodríguez-Farré, 1997; Soler-Martín et al., 2007; Wilkerson et al., 2018). IDPN (>98%, TCI Europe, Zwijndrecht, Belgium) was administered i.p. in 2 ml/kg of saline. The first lot of animals was used in an experiment in which rats were administered a single dose of 0 (Control group), 400 (IDPN 400), 600 (IDPN 600), or 1000 (IDPN 1000) mg/kg of IDPN (n=6 / group). Behavioral and scanning electron microscopy data from this experiment have been published elsewhere (Martins-Lopes et al., 2019). For the present study, HC counts were obtained by immunofluorescent analysis from the second ear of these animals. In the second experiment, animals received 0, 450 (IDPN 450), 500 (IDPN 500), 550 (IDPN 550), or 600 mg/kg of IDPN (n=5/group). In the third experiment, rats were administered with 0, 150 (IDPN 3×150), 175 (IDPN 3×175), or 200 (IDPN 3×200) mg/kg·day of IDPN for 3 consecutive days (n=6/group). Administration of IDPN in three doses over three days has been reported to cause graded ototoxicity with limited systemic toxicity (Crofton and Knight, 1991). The animals were evaluated for vestibular function before exposure and at regular time-points after exposure for 13 weeks (experiment 1) or 4 weeks (experiments 2 and 3). The time-course data from experiment 1 (Martins-Lopes et al., 2019) indicated a stability in the reflex deficits between 4 and 13 weeks. Also, previous histological observations indicate that additional HC loss is unlikely to occur beyond 3 weeks after IDPN administration (Llorens and Demêmes, 1994). The tail-lift reflex was assessed in the three lots of animals and the air-righting reflex was assessed only in experiments 1 and 3.

At the end of the experimental periods, rats were given an overdose of anesthesia and decapitated for the purpose of isolating the vestibular sensory epithelia. The temporal bones were immersed in cold fixative and immediately dissected under a fume hood. The first inner ear of animals in the first experiment was used for scanning electron microscopy as published elsewhere (Martins-Lopes et al., 2019). The other inner ears were collected for the present study.

### 2.2. Assessment of vestibular reflexes

The tail-lift reflex and the air-righting reflex were assessed using high-speed video recording as described (Martins-Lopes et al, 2019; Maroto et al., 2021). Briefly, in the tail-lift reflex test, the rat is grasped by the base of the tail, gently lifted to approximately 40 cm, and then lowered down to the starting point (see Supplementary movies A and B for the tail-lift reflex in control and vestibular deficient rats). To facilitate image tracking, a white marble is placed in the back of the neck of the rat with a rubber band collar and the test is done in front of a red background. In the air-righting reflex test, the experimenter holds the rat with the two hands in a supine position at approximately 40 cm of height and suddenly releases it to fall on a foam cushion (see Supplementary movies C and D for the air-righting reflex in control and vestibular deficient rats). The reflex behaviors were recorded at 240 frames per second with a Casio Exilim ZR700 or a GoPro Hero 5 camera. Using the free software Kinovea (www.kinovea.org) for video tracking, we obtained the coordinates of the nose, back of the neck and base of the tail at a 1/240 s frequency during the tail-lift maneuver, and the time from the release of the animal until it fully righted its head in the air-righting reflex test. A script in R programming language was then used to calculate the minimum angle from the nose-neck-tail coordinates during the tail-lift test (Martins-Lopes et al., 2019).

### 2.3. Immunohistochemistry

The following primary antibodies were used: rabbit anti-Myosin VIIa (Myo7a) from Proteus Biosciences (cat. # 25-6790), rabbit anti-tenascin from Millipore (cat. # AB19013), rabbit anti-oncomodulin from Swant (cat. # OMG4), guinea pig anti-calretinin from Synaptic Systems (cat. # 214-104), mouse anti-Myo7a (clone 138-1-s, IgG1, supernatant) from Developmental Studies Hybridoma Bank, mouse anti-CtBP2/Ribeye (clone 16/CtBP2, IgG1, cat. # 612044) from BD Biosciences, mouse anti-contactin-associated protein (Caspr1) (clone K65/35, IgG1, cat. # 75-001) and anti-PSD95 (clone K28/43, IgG2a, cat# 75-028) from Neuromab. We used the following secondary antibodies conjugated with Alexa Fluor fluorochromes: 488 goat anti-guinea-pig IgG H+L (catalog #A11073, Invitrogen/ThermoFisher), 555 donkey anti-rabbit IgGs H+L (#A-31572), 555 goat anti-mouse IgG2a (#A21137), 647 goat anti-mouse IgG1 (#A21240) and 654 donkey anti-mouse IgG H+L (catalog #A-21202). We also used the DyLight 405 donkey anti-rabbit IgG H+L (catalog #711-475-152, Jackson ImmunoResearch). The nuclear stain 4□,6-diamidino-2-phenylindole (DAPI) was obtained from Sigma.

Vestibular epithelia were dissected in cold 4 % freshly depolymerized paraformaldehyde in phosphate buffered saline (PBS). After dissection, the tissues were fixed for 1 h in the same fixative, transferred to a cryoprotective solution (34.5% glycerol, 30% ethylene glycol, 20% PBS, 15.5% distilled water) and stored at −20°C until further processing. For immunolabelling, we followed the protocol by Lysakowski et al. (2011). For most animals, one horizontal crista, one utricle and one saccule were processed. Briefly, samples were placed at room temperature, rinsed, permeabilized with 4% Triton X-100 in PBS (1 h) and blocked with 1% of fish gelatin in PBS containing 0,5% Triton X-100 (1 h). Primary antibodies were incubated in 0.1% Triton X-100 and 1% fish gelatin in PBS for 24h at 4°C. Samples from all animals were immunolabelled with rabbit anti-Myo7a (1/400), mouse anti-Caspr1 (1/400) and guinea-pig anti-calretinin (1/500). Samples from the second ear of selected animals were also immunolabelled with mouse anti-Myo7a (1/100), rabbit anti-tenascin (1/200), rabbit anti-oncomodulin (1/400), and guinea-pig anti-calretinin (1/500). To assess synaptic puncta, we combined the rabbit anti-Myo7a and the guinea pig anti-calretinin with the mouse IgG1 anti-CtBP2/Ribeye and mouse IgG2a anti-PSD95 antibodies to label one anterior or posterior crista. Available data suggest that all three cristae are similarly affected by IDPN (Llorens et al., 1993; Llorens and Demêmes, 1994; Seoane et al., 2001a). Secondary antibodies were incubated in the dark using the same buffer and conditions used with the primary antibodies. Specimens were thoroughly rinsed with PBS after each incubation period. When only three primary antibodies had been used, an incubation of 15 min with DAPI (1/1000) in PBS was intercalated among the final rinses. To assess synaptic contacts, the labelled specimens were included in a bloc of gelatin/albumin as described (Sedó-Cabezón et al., 2015) and sectioned at 40 μm in a vibrating blade microtome (Leica VT1000S). The whole epithelia or the epithelial sections were finally mounted in Mowiol medium.

### 2.4. Confocal microscopy, HC counts and synaptic counts

Vestibular epithelia were visualized in a Zeiss LSM880 spectral confocal microscope using a 63× (NA: 1.4) objective. Z-stacks of optical sections 0,5 μm thick (for HC counts) or 0,3 μm thick (for synaptic counts) were obtained spanning the whole sensory epithelium. HC counts were obtained for images from one central/striolar region and one (crista and saccule) or two (utricle) peripheral regions. The imaged regions were selected by cytoarchitectonic criteria when available (presence of calretinin+ calyces or oncomodulin+ HCs) or by approximate localization (as shown in Fig. 1B) when these had been lost due to the IDPN treatment. In the utricle, separate counts were obtained from the peripheral regions at the external (lateral) and internal (medial) sides to the striola. The obtained thick 3D images were processed with the blend option of the Imaris image processing software (Bitplane), and the resulting filtered images used for cell counts using the ImageJ software (National Institute of Mental Health, Bethesda, Maryland, USA).

The first immunolabeling combination was used to obtain estimates of the numbers of HCs, HCIs and HCII. First, all HCs were identified by the cytoplasmic labelling of the anti-Myo7a antibody (Hasson et al., 1997). Second, the Caspr1 label of the calyceal junctions of the calyces was used to count HCIs (Sousa et al., 2009; Lysakowski et al., 2011). As argued in the Discussion section below, this label provided a reliable identification of HCIs in the present model, in which HCs either degenerate or survive the toxic insult. Third, we counted cells with overlapping calretinin and Myo7a cytoplasmic label as HCIIs (Dechesne et al., 1991). The calretinin label of calyx-only afferents in the central areas of the receptors is clearly distinguished from that of HCIIs (Desmadryl and Dechesne, 1992). Cell counts per stack were obtained manually because we failed to obtain reliable counts using the segmentation functions of the programs used for image processing. Cells transected by the borders of the frame were not counted. As the three counts were obtained using separate images obtained by different combinations of color channels from the same stack, sums of HCI and HCII counts were not identical to all HC counts. In selected animals from the second and third lots, a second combination was used that included the anti-oncomodulin antibody as marker to prominently delineate the central zone of the receptors (Hoffman et al., 2018). With this combination, HCI from peripheral zones of the receptors were identified by the tenascin expression at the calyceal junctions (Lysakowski et al., 2011). In the synaptic study, we counted numbers of pre-synaptic (ribeye) and post-synaptic (PSD-95) puncta on HCII defined by the colabelling with the anti-Myo7a and anti-calretinin antibodies (Sedó-Cabezón et al., 2015; Greguske et al., 2019).

### 2.5. Statistics

Data are shown as mean +/- SE per dose group. Group comparisons were performed using one-way ANOVA followed by Duncan’s post-hoc tests. To evaluate the relationship between histology counts and behavioral data, two limit values were set to define tail-lift angles and air-righting times corresponding to normal, altered and absent vestibular function, as explained in the Results section. Then, cell counts from the three groups of animals defined by these limit values of angle or time were compared by Kruskal-Wallis ANOVA followed by pairwise Dunn-Bonferroni pots-hoc comparisons. Because IDPN1000 animals showed a complete loss of HCs in all epithelia and zones, this group was excluded from these analyses. Student’s t-test was used to compare synaptic count data between samples from control and treated animals. All analyses were done with the IBM SPSS 25 program package.

## 3. RESULTS

### 3.1. Effects of IDPN ototoxicity on anti-gravity reflexes

Rats exposed to IDPN showed effects on body weight, spontaneous behavior, tail-lift reflex, and air-righting reflex in accordance with previous observations (Crofton and Knight, 1991; Llorens et al., 1993; Martins-Lopes et al., 2019). Body weight loss was transient, and animals returned to weight increase by one week after IDPN administration. The first experiment included rats exposed to 0, 400, 600 and 1000 mg/kg of IDPN and recorded dose- and time-dependent effects on the vestibular reflexes, as previously published (Martins-Lopes et al., 2019). The findings for the tail-lift reflex were confirmed and extended in experiments 2, in which rats received 0, 400, 450, 500, 550 or 600 mg/kg of IDPN, and 3, in which animals were dosed over 3 consecutive days at 0, 3 x 150, 3 x 175, or 3 x 200 mg/kg·day (Supplementary Fig. S1). To obtain a single measure of the final effect of the ototoxic treatment on this reflex, we averaged the tail-lift angles from day 21 to the end of the experiment at days 28 (experiments 2 and 3) or 91 (experiment 1), as shown in Fig. 2A-C. The ventral curling of the vestibular-deficient rats results in low angles (Fig. 2D). ANOVA analysis of the averaged tail-lift angles for experiments 1, 2, and 3 resulted, respectively, in F[3, 20] = 189.3 (p = 0.000), F[4, 20] = 7.296 (p = 0.001), and F [3, 17] = 13.44 (p = 0.000).

**Figure 2.**
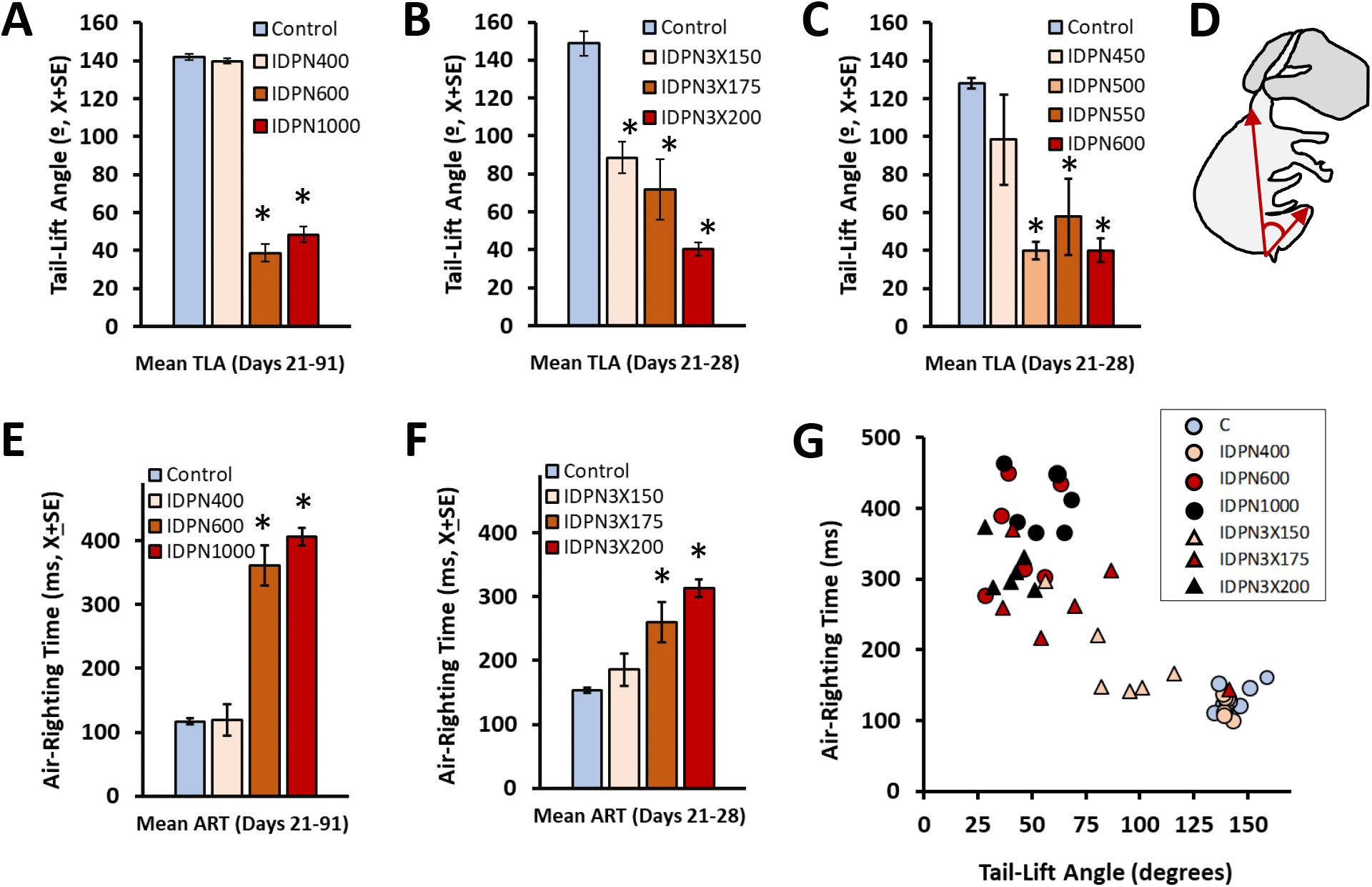
Effects of the vestibular toxicity of IDPN on the tail-lift and air-righting reflexes. (A-C) Mean ± SE minimum nose-neck-base of the tail angles displayed by the rats when lifted by the tail and lowered back.). (D) Schematic drawing of the tail-lift angle as obtained from a rat suffering a deep loss of vestibular function. (E-F) Mean ± SE air-righting times displayed by the rats when dropped in supine position from approximately 40 cm above a foam cushion. The data are average values from days 21 to 91 after exposure (A and E, 0 - 1000 mg/kg, one dose, n=6/group) or days 21 to 28 after exposure (B and F, 0 - 200 mg/kg·day, for 3 consecutive days, n=6/group; and C, 0 - 600 mg/kg, one dose, n=5/group). (G) Relationship between individual values of the tail-lift angle and the time to right in the animals whose group values are shown in panels A, B, E and F. *: p<0.05, significantly different from control group, Duncan’s test after significant ANOVA.

In agreement with the previous data on the effects of IDPN on the air-righting reflex (experiment 1, Martins-Lopes et al., 2019), a 3-day exposure to IDPN (experiment 3) was found to cause an impairment in performance resulting in dose- and time-dependent increases in air-righting times (Supplementary Fig. 2). Figure 2E-F show the average air-righting times from day 21 to the end of the experiment for experiments 1 and 3. ANOVA analysis of these averaged times resulted in F[3, 20] = 76.15 (p = 0.000), and F [3, 17] = 7.705 (p = 0.002), respectively.

Figure 2G shows the relationship between the tail-lift angles and the air-righting times obtained in individual animals. All control animals had average tail-lift angles above 120 degrees and average air-righting times below 170 ms, so these values were selected as limits to differentiate normal versus abnormal average responses in subsequent analyses. The high IDPN1000 dose group showed average tail-lift angles below 70 degrees and air-righting times above 350 ms. These values defined a second set of limits for responses corresponding to an absence of vestibular input, because the IDPN1000 animals displayed a complete loss of HCs (see below). The dose-dependent effects of IDPN on each of the two reflexes was different and tail-lift angles decreased at doses below those at which air-righting times increased. Thus, four of the six animals in the IDPN3X150 group had angles in the 70-120 range, below the normal range, while their air-righting times were in the normal range (below 170 ms). Taking all animals into account, only 6 out of 69 rats had tail-lift angles between 70 and 120 degrees, so a majority of the animals in the study could be classified as having either a normal or an absent reflex, and less than 10 % showed a smaller than normal but present response. In the air-righting reflex, we found 14 out of 44 rats with air-righting times between 170 and 350 ms, that is, a 30 % of animals with a slower than normal but present reflex.

### 3.2. Dose-, epithelial zone-, and type-dependent loss of HCs after IDPN exposure

The vestibular sensory epithelia of control rats (Fig. 3–4) showed the expected high density of HCs (Myo7a+), including HCI (identified by the Caspr+ calyx around the basolateral end of the Myo7a+ cytoplasm) and HCII (with cytoplasmic calretinin and Myo7a co-labelling and no calyx). In the macular striola and central crista zones, most of the HCIs were contacted by calyx-only afferent terminals expressing calretinin. The shape of this calretinin label was very different from, and therefore not confused with, the cytoplasmic label of calretinin in HCII, which co-localizes with the Myo7a label.

**Figure 3.**
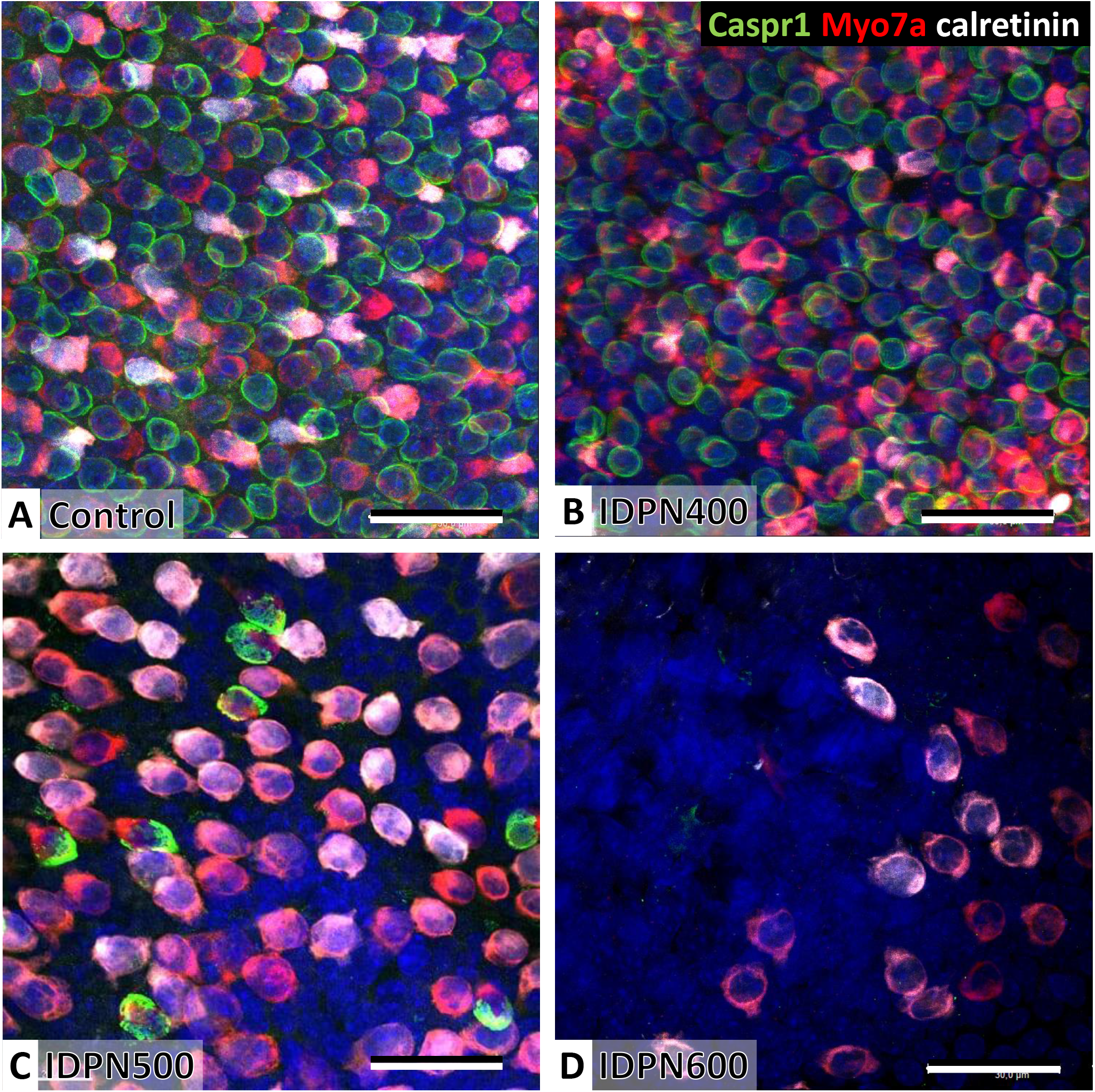
Effect of ototoxic exposure on the density of sensory hair cells (HCs) in the vestibular epithelia. The example images correspond to the medial part of the utricle, peripheral to the striola. Type I HCs (HCI) are revealed by the Caspr+ label on the inner membrane of the calyx (green). Type II HCs (HCII) show no Caspr+ label but are Myo7a+ (red) and calretinin+ (white). Nuclei were labeled with DAPI (blue). Images shown here are stack projections optimized to provide an overall sense of HC density, not an accurate identification of the cell types. Cells that seem to be Myo7a+, Caspr- and calretinin-may be HCII that express calretinin at low levels or HCI with their calyceal junction in planes not included in the image, and Caspr1+ calyces with no visible Myo7a+ are HCI cells with suboptimal red mark in this particular image. Also, HC superposition may generate apparent calretinin label in Caspr1+ cells. For cell counts, color channels were split as shown in Fig. 4. (A) Normal density of cells in a control rat. (B) Control-like appearance was found in most rats receiving the lowest dose of IDPN (400 mg/kg). (C) Overt loss of HCI and HCII in an utricle of a rat dosed with 500 mg/kg of IDPN. (D) Complete loss of HCI and nearly complete loss of HCII in a rat dosed with 600 mg/kg of IDPN. Scale bars = 30 μm.

**Figure 4.**
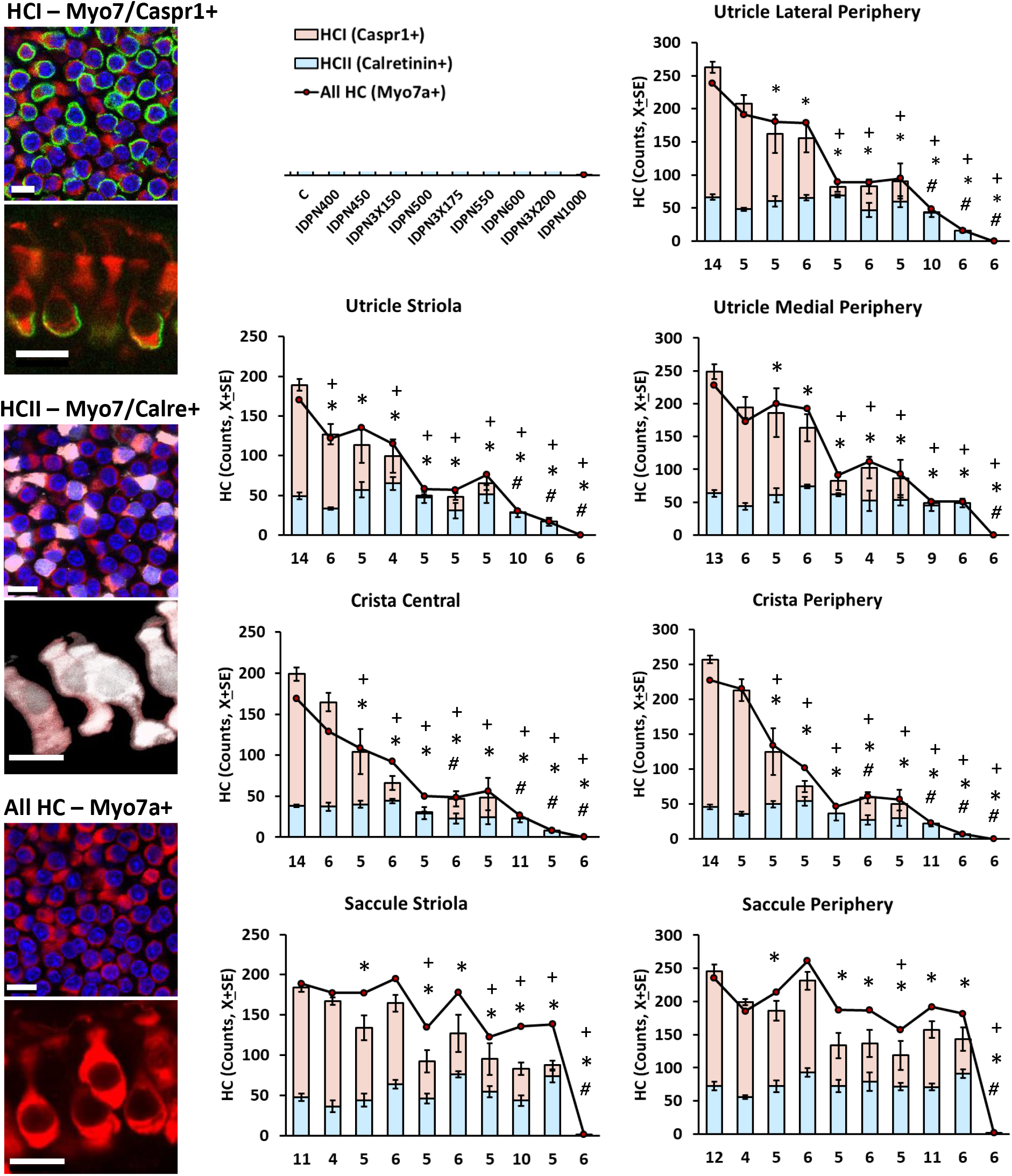
Differences in the loss of type I hair cells (HCI), type II hair cells (HCII), and all hair cells (HC) after exposure to the ototoxic compound, IDPN, as a function of the dose (400 to 1000 mg/kg, see legend), zone (central/striola vs periphery) and end-organ (utricle, crista, and saccule). The images in the left column show the use of the Caspr1 (green) and Myo7a (red) label to identify HCI, use of the Calretinin (white) and Myo7a (red) label to identify HCII and the use of Myo7a (red) label to identify all HCs. Nuclei in blue were labelled with DAPI. Scale bars = 10 μm. Bar graphs show superposed HCI and HCII counts (X±SE) according to end-organ, region within it and IDPN dose. The line show mean All HC counts. Error bars for All HCs are not included for clarity. The sum HCI+HCII and All HCs are not identical in all cases because the values correspond to counts obtained from separate images. Numbers below the bars indicate numbers of animals. #, *, +: p<0.05, significantly different from control group for HCII, HCI, and All HC, respectively, by Duncan’s test after significant ANOVA. Note that loss of HCI starts at low doses of IDPN, while these low doses have no effect on HCII counts, particularly in the saccule.

IDPN caused a dose-dependent loss of HCs as observed in previous studies (Llorens et al., 1993; Llorens and Demêmes, 1994; Martins-Lopes et al., 2019). The time course of the HC degeneration induced by acute or sub-acute IDPN has been well characterized, occurs mostly within one week after dosing, is completed by three weeks and does not progress after this time (Llorens et al., 1993; Llorens and Demêmes, 1994; Seoane et al., 2001a). Therefore, the data collected represent the final outcome of the toxic lesion. While most of the samples from IDPN400 animals showed a control-like overall appearance (Fig 3B), vestibular epithelia from IDPN1000 animals showed no HCs, except for a few sparse HCII remaining in the saccule of 2 of the 6 animals. Therefore, motor responses recorded in the IDPN1000 animals correspond to those independent of vestibular input. Other dose groups showed intermediate degrees of HC loss that varied as a function of HC type, receptor, and zone (Fig. 3C, D). For a quantitative analysis of these differences, counts of HCI (Caspr+), HCII (calretinin+) and all HCs (Myo7a+) were obtained from all the animals in the three behavioral experiments in seven different fields of observation: striola of the utricle, external extra-striola (lateral periphery) of the utricle, internal extra-striola (medial periphery) of the utricle, central crista, peripheral crista, striola of the saccule, and periphery of the saccule. The histological data of animals from the three behavioral experiments were analyzed as a single experiment as shown in compact format in figure 4 and type-by-type in Supplementary Figures S3-S5. Significant effects of IDPN were observed in all 21 ANOVA comparisons (all F[9, 53-59] > 7.98, p’s = 0.000), but different dose response relationships were evident in different zones and cell type. For instance, a total loss of HCI was recorded in the central and peripheral crista of IDPN600 animals, while comparison of this group with the Control group in the saccular periphery revealed a 50 % of survival of HCIs and no significant effect on HCII counts (Fig. 4). As another example, the fact that HCII counts in the saccular periphery were only reduced after the highest dose (1000 mg/kg) of IDPN clearly differed from the significant 33% decrease in HCI recorded already at the lowest one (400 mg/kg) in the striola of the utricle (Fig. 4). Figure 4 also shows that we recorded counts of all HCs somewhat lower than the sum of HCI and HCII counts in the crista and utricles of Control rats and in the crista center and utricle peripheries of IDPN400 rats. In contrast, counts of all HCs higher than HCI+HCII counts were recorded in the saccular striola and periphery of several treatment groups, as well as in the crista of IDPN3×150 rats.

The numbers of HCs of each type and in each epithelial zone were estimated using a second antibody combination with samples from the second ear of the rats in the second and third experiment. In this combination, oncomodulin delineates the central/striola zone of the receptors. The data gathered (see supplementary material Fig. S6-S8), provided a confirmatory replicate of the different dose-response relationships that characterize different zones and cell types.

### 3.3. Relationship between HC loss and deficits in anti-gravity reflexes

As a first approximation to identify the cellular basis of the tail-lift reflex and the air-righting reflex, we plotted the HC counts by type of cell, epithelium and zone against the quantitative measures of the reflexes. As shown in Fig. 5–10, the expected overall basic pattern was recorded: different doses of the ototoxic compound caused varying degrees of HC loss and in parallel decreases in tail-lift angles and increases in air-righting times were recorded. However, the relationship between the cell count and behavioral effect varied as a function of the cell type, end-organ and epithelial zone considered. The IDPN1000 rats were excluded for statistical analyses of these relationships because they showed no HC remaining in any of the vestibular epithelia.

**Figure 5.**
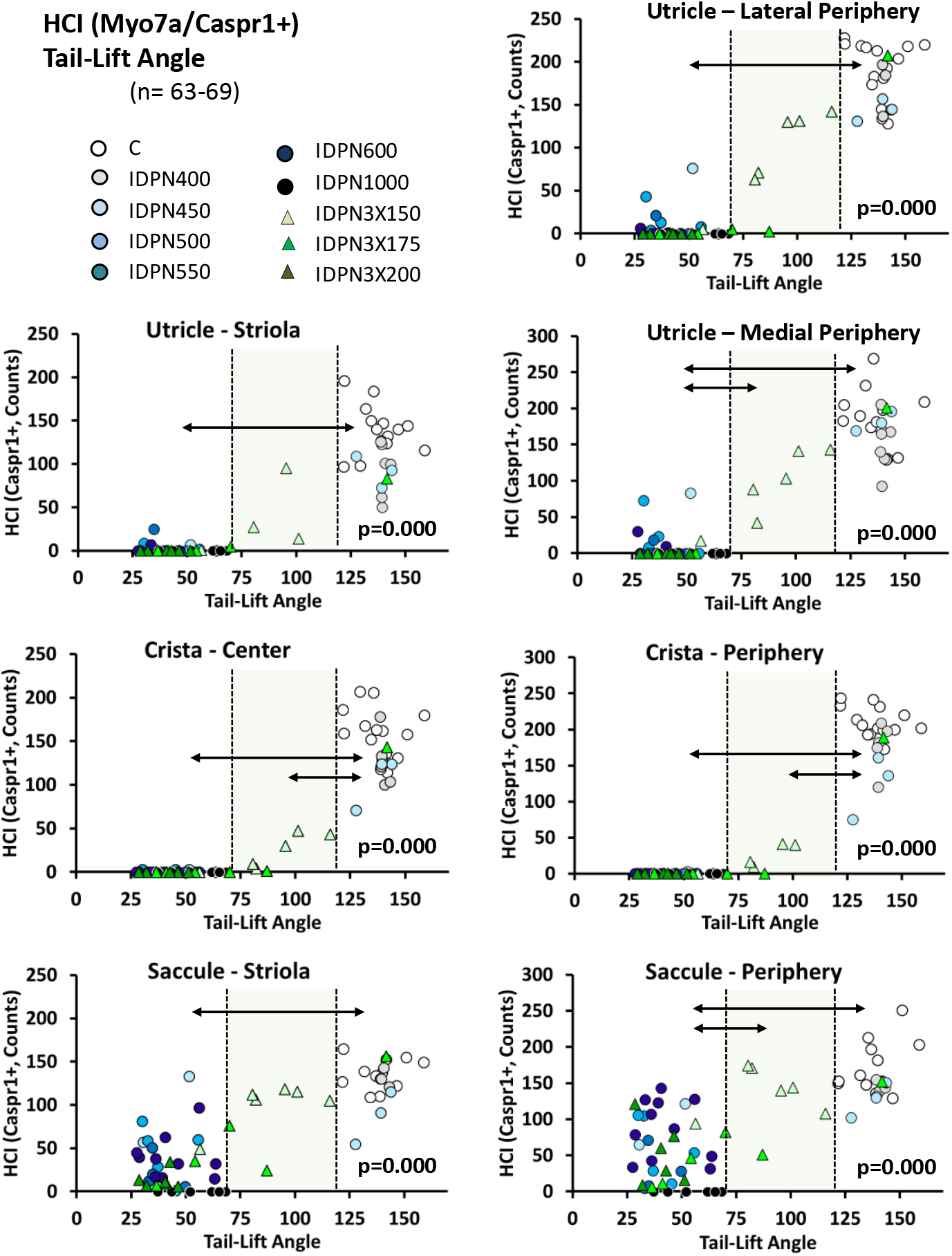
Relationship between HCI loss and tail-lift angle decrease after exposure to a variety of doses of the ototoxic compound, IDPN, as a function of the zone (central/striola vs periphery) and end-organ (utricle, crista and saccule). Individual data shown here correspond to those shown as group means in Fig. 2A-C (angles) and Fig. 4 (HCI counts). On each panel, the vertical dashed lines indicate the limit of normal angles (120 degrees, defined by the Control group) and angles in complete absence of vestibular function (70 degrees, defined by the IDPN1000 group). These limit values classified rats into three groups with high, medium and low angles. The p value in each panel indicates statistical significance in HCI counts among these three groups of rats, after exclusion of the IDPN1000 animals. Double-headed arrows denote significance (p<0.05) of the post-hoc pair-wise comparisons.

**Figure 6.**
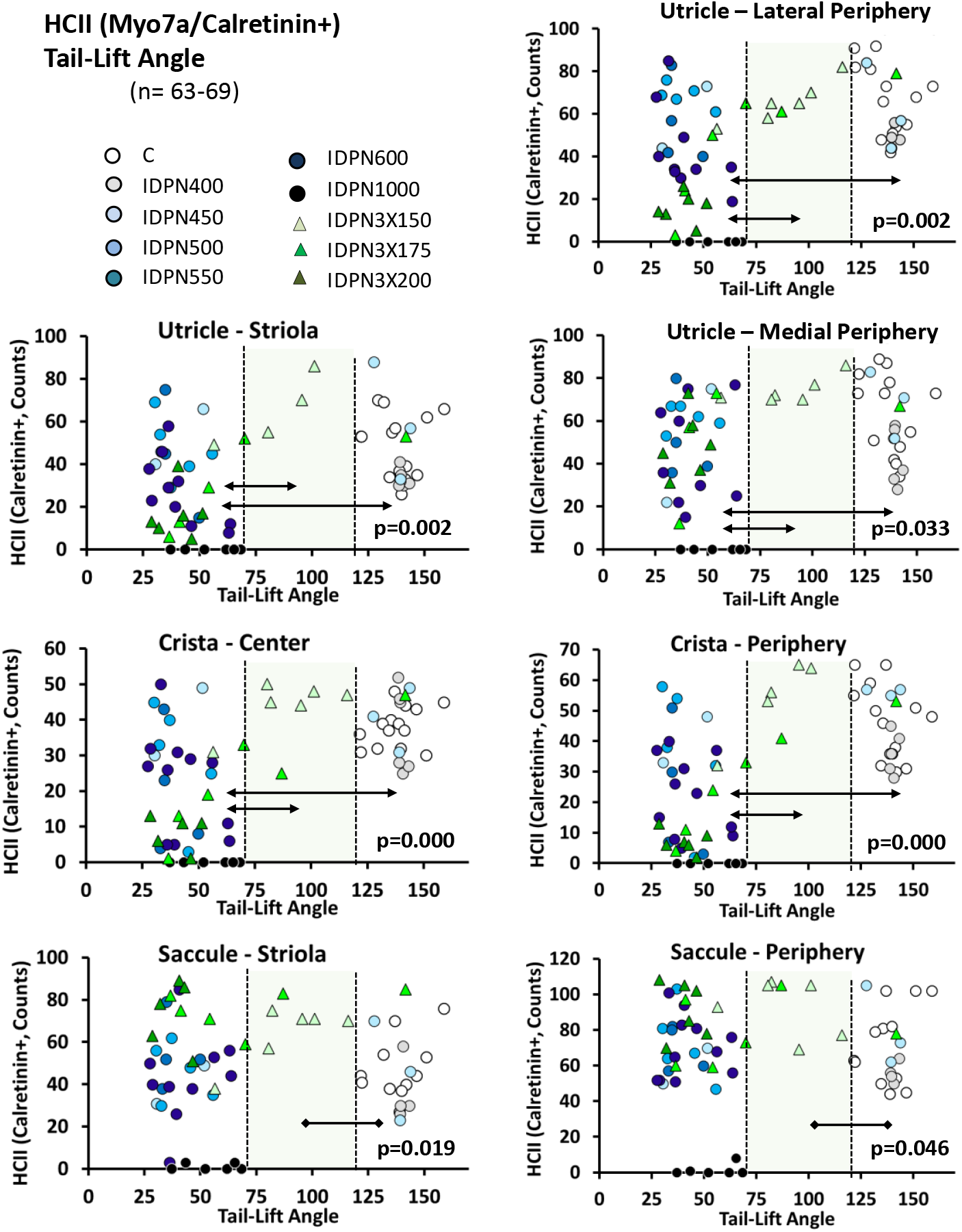
Relationship between HCII loss and tail-lift angle decrease after exposure to a variety of doses of the ototoxic compound, IDPN, as a function of the zone (central/striola vs periphery) and end-organ (utricle, crista and saccule). Individual data shown here correspond to those shown as group means in Fig. 2A-C (angles) and Fig. 4 (HCII counts). On each panel, the vertical dashed lines indicate the limits of normal angles (120 degrees, defined by the Control group) and angles in complete absence of vestibular function (70 degrees, defined by the IDPN1000 group). These limit values classified rats into three groups with high, medium and low angles. The p value in each panel indicates statistical significance in HCII counts among these three groups of rats, after exclusion of the IDPN1000 animals. Double-headed arrows denote significance (p<0.05) of the post-hoc pair-wise comparisons. In the saccule periphery and striola, HCII counts in medium angle animals were higher than in high angle animals.

**Figure 7.**
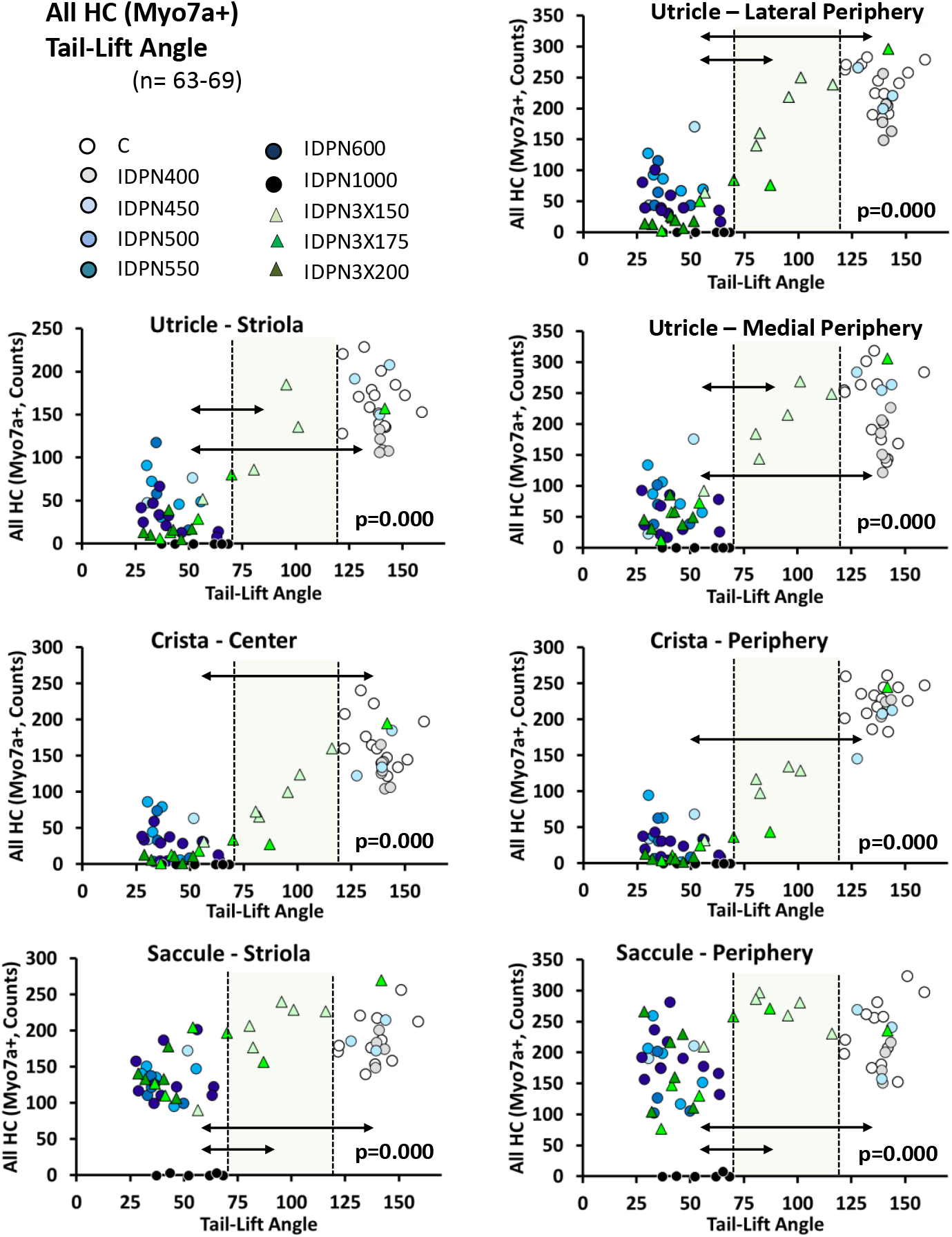
Relationship between all HC loss and tail-lift angle decrease after exposure to a variety of doses of the ototoxic compound, IDPN, as a function of the zone (central/striola vs periphery) and end-organ (utricle, crista and saccule). Individual data shown here correspond to those shown as group means in Fig. 2A-C (angles) and Fig. 4 (all HC counts). On each panel, the vertical dashed lines indicate the limits of normal angles (120 degrees, defined by the Control group) and angles in complete absence of vestibular function (70 degrees, defined by the IDPN1000 group). These limit values classified rats into three groups with high, medium and low angles. The p value in each panel indicates statistical significance among these three groups of rats, after exclusion of the IDPN1000 animals. Double-headed arrows denote significance (p<0.05) of the post-hoc pair-wise comparisons.

**Figure 8.**
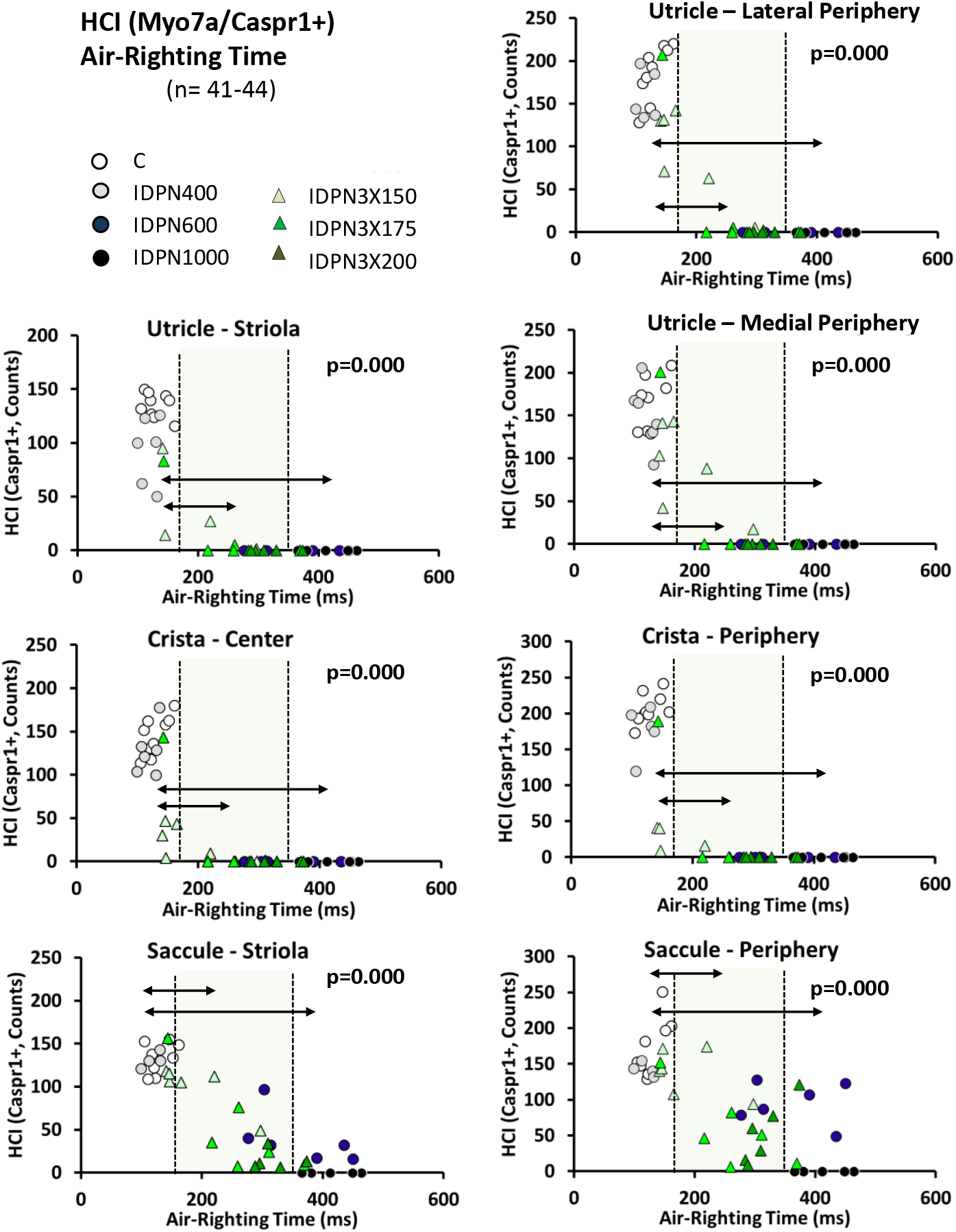
Relationship between HCI loss and air-righting time increase after exposure to a variety of doses of the ototoxic compound, IDPN, as a function of the zone (central/striola vs periphery) and end-organ (utricle, crista and saccule). Individual data shown here correspond to those shown as group means in Fig. 2E-F (times) and Fig. 4 (HCI counts). On each panel, the vertical dashed lines indicate the limits of normal times (170 ms, defined by the Control group) and times in complete absence of vestibular function (350 ms, defined by the IDPN1000 group). These limit values classified rats into three groups with low, medium and high times. The p value in each panel indicates statistical significance among these three groups of rats, after exclusion of the IDPN1000 animals. Double-headed arrows denote significance (p<0.05) of the post-hoc pair-wise comparisons.

**Figure 9.**
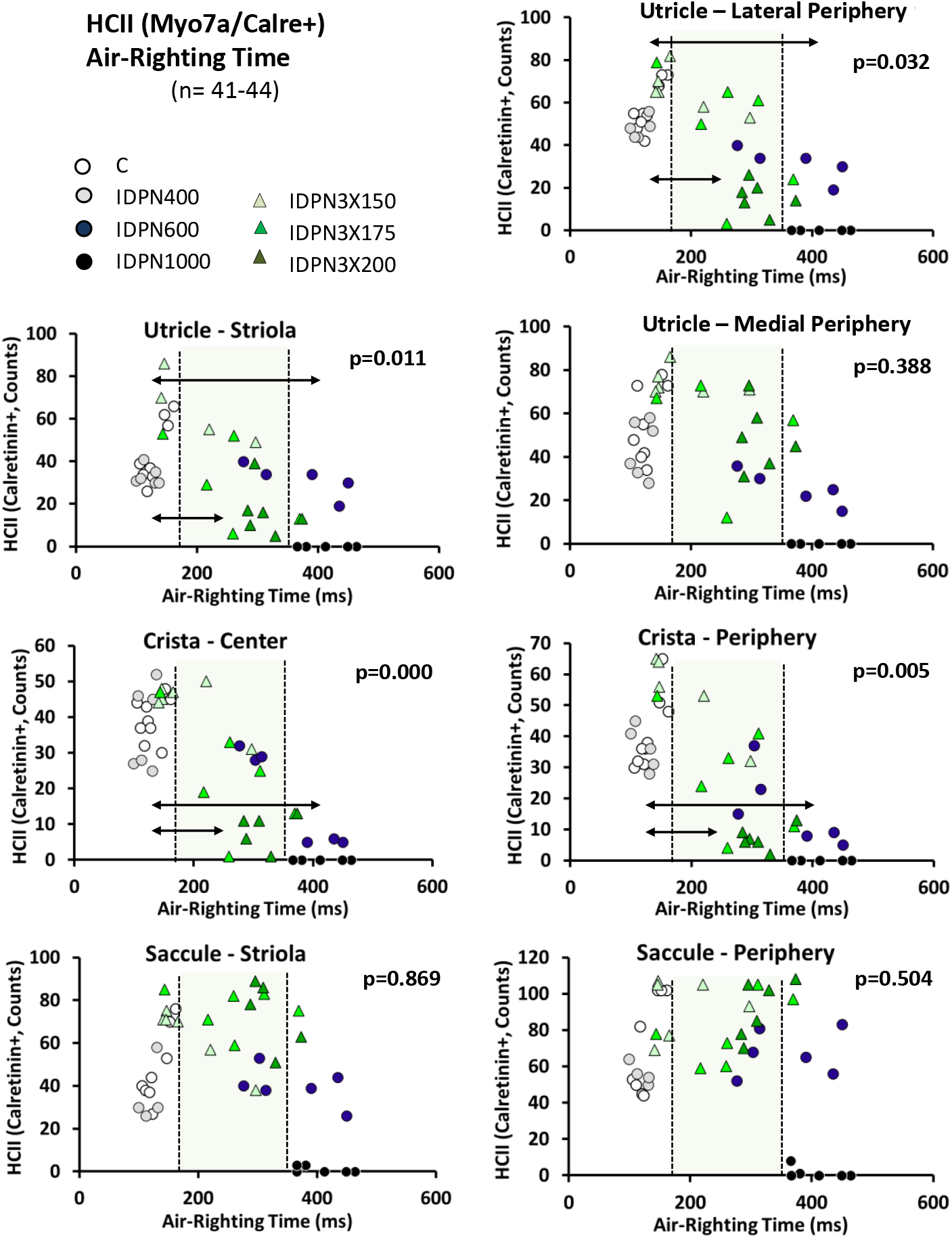
Relationship between HCII loss and air-righting time increase after exposure to a variety of doses of the ototoxic compound, IDPN, as a function of the zone (central/striola vs periphery) and end-organ (utricle, crista and saccule). Individual data shown here correspond to those shown as group means in Fig. 2E-F (times) and Fig. 4 (HCII counts). On each panel, the vertical dashed lines indicate the limits of normal times (170 ms, defined by the Control group) and times in complete absence of vestibular function (350 ms, defined by the IDPN1000 group). These limit values classified rats into three groups with low, medium and high times. The p value in each panel indicates statistical significance among these three groups of rats, after exclusion of the IDPN1000 animals. Double-headed arrows denote significance (p<0.05) of the post-hoc pair-wise comparisons.

**Figure 10.**
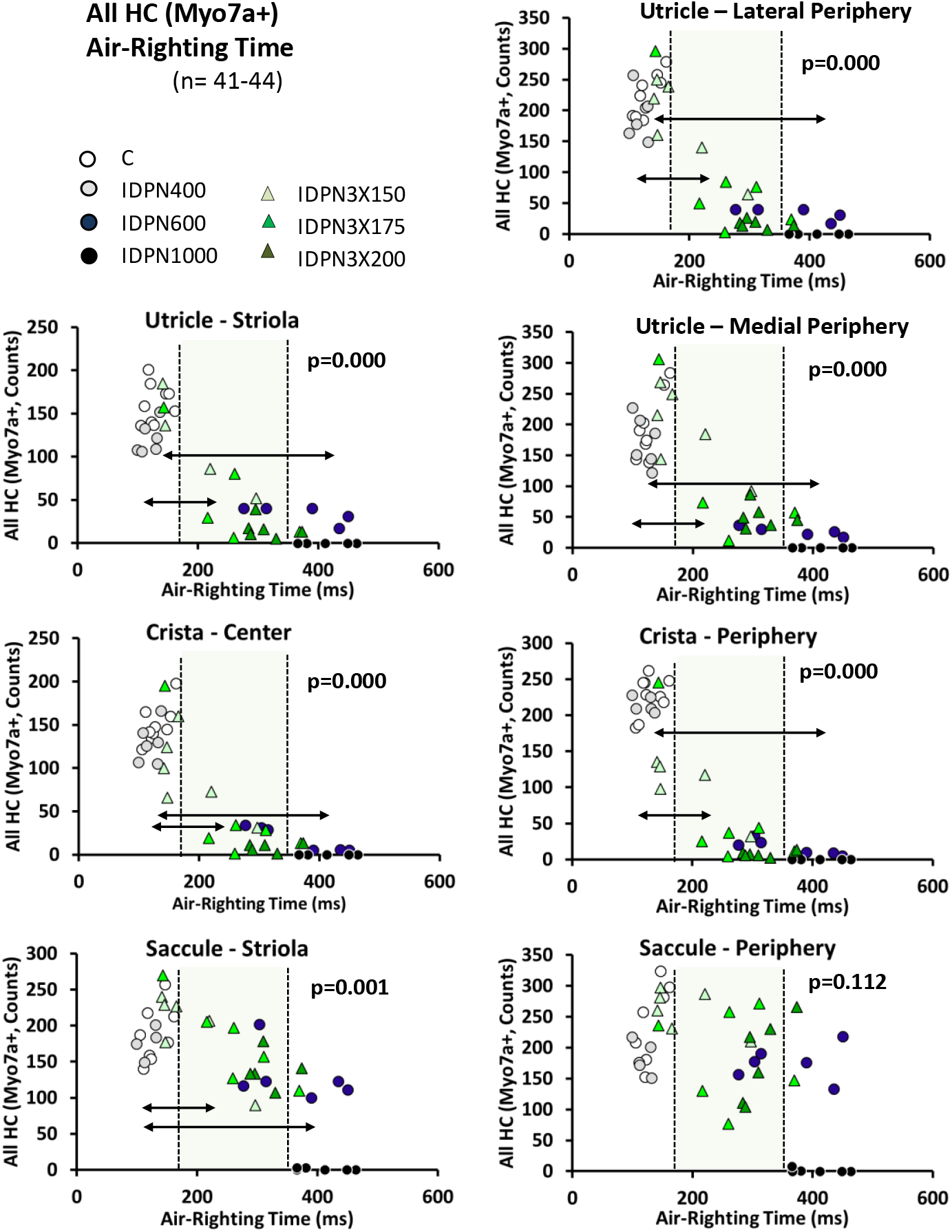
Relationship between all HC loss and air-righting time increase after exposure to a variety of doses of the ototoxic compound, IDPN, as a function of the zone (central/striola vs periphery) and end-organ (utricle, crista and saccule). Individual data shown here correspond to those shown as group means in Fig. 2E-F (times) and Fig. 4 (all HC counts). On each panel, vertical dashed lines indicate the limits of normal times (170 ms, defined by the Control group) and times in complete absence of vestibular function (350 ms, defined by the IDPN1000 group). These limit values classified rats into three groups with low, medium and high times. The p value in each panel indicates statistical significance among these three groups of rats, after exclusion of the IDPN1000 animals. Double-headed arrows denote significance (p<0.05) of the post-hoc pair-wise comparisons.

As shown in Fig. 5, low tail-lift angles associated with low numbers of HCI counts in all epithelia and zones. In all cases, statistically significant differences were found among HCI numbers from animals showing high (above 120 degrees, normal), medium (between 120 and 70 degrees, reduced vestibular function) or low (below this limit value, absent function) angles. In pair-wise comparisons, significant differences were found between animals with high and low angles in all epithelia and zones. HCI counts in the crista center and periphery were significantly lower in the medium than in the high angle animals, whereas significant differences were recorded between the medium and low angle animals in the utricle medial periphery and the saccule periphery. In contrast, a significant decrease in HCII counts (Fig. 6) was recorded in all crista and utricle zones to occur between medium and low angle animals, not between high and medium angle animals. In the saccule, an apparent increase in HCII occurred in medium angle compared to high angle animals, but no difference was found between high and low angle animals. With the exception of the IDPN1000 animals, which had been excluded from the statistical analyses, many treated animals showed a complete loss of the tail-lift extension reflex while still retaining a control-like density of HCII in the saccule. When all (Myo7a+) HCs were considered, significant differences were recorded in all epithelia and zones between animals with angles above the 120 limit and those with angles below the 70 degrees limit. No significant differences were recorded between medium and high angle animals, while significant differences occurred in the utricle and saccule regions between medium and low angle animals.

The relationship between air-righting times and HCI counts are shown in Fig 8. For all epithelia and zones, the groups of animals with times greater than the 170 and 350 ms limit values had significantly lower numbers of HCIs than the rats showing times within the normal range. However, the group of animals with air-righting times within the normal range included animals with frankly reduced numbers of HCI in the crista and utricle. For HCII (Fig. 9), statistically significant differences were found between low and medium and between low and high times in the crista center, crista periphery, utricle striola and utricle lateral periphery, but not for utricle medial periphery or saccular striola and periphery. Significant count differences were found for all (Myo7a+) HCs (Fig. 10) between low and medium and between low and high times in all epithelia and zones except for the saccular periphery.

The relationships between the behavioral and histological data were also examined using the second series of immunochemical labels. These labels offered a more precise localization of the striola/center regions of the epithelia, but a smaller number of samples were available for analysis. As shown in Supplementary Fig. S9 to S14, the results obtained with this second combination of antibodies were similar to those obtained with the first combination.

### 3.4. Synaptic puncta numbers in surviving HCII

The comparisons of cell numbers shown above clearly indicated that HCI are more sensitive to IDPN toxicity than HCII. However, surviving cells may have partially or totally lost function (Hirvonen et al., 2005). Recent research has shown that synaptic uncoupling is one of the causes of functional loss before HC loss becomes evident (Sedó-Cabezón et al., 2015; Sultemeier and Hoffman, 2017; Cassel et al., 2019; Greguske et al., 2019). To evaluate the possibility that surviving HCII were suffering synaptic uncoupling, we assessed pre- and post-synaptic puncta numbers in cristae of animals selected to be representative of the different degrees of behavioral dysfunction. The results obtained (Fig. 11) did not support that surviving HCII were suffering synaptic uncoupling. Thus, the number of pre-synaptic Ribeye puncta and of the post-synaptic PSD95 puncta were similar in HCII of control and IDPN-treated animals. Student’s test t values were 0.138 for ribeye puncta and 0.403 for PSD95 puncta, both p > 0.05 (20 d.o.f.). Also, no significant differences were found for numbers of ribeye or PSD95 puncta per HCII cell among groups of rats with high, medium, or low tail-lift angles, or among rats with low, medium, or high air-righting times.

**Figure 11.**
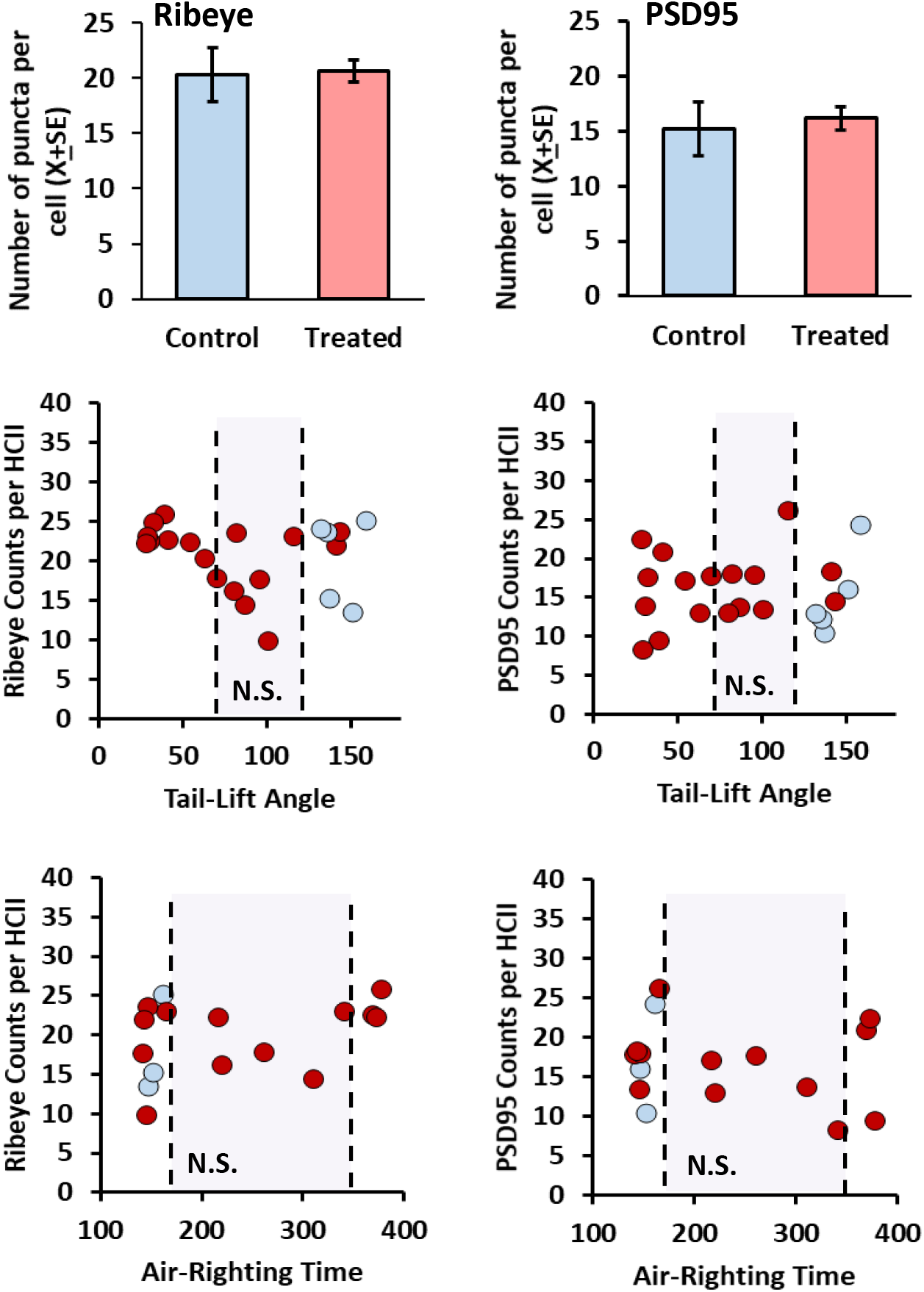
Counts of pre-synaptic (Ribeye+) puncta and post-synaptic (PSD95+) puncta in HCII of control rats (n=5) and rats with graded vestibular lesions (n=17), selected to cover the spectrum of values of tail-lift angles and air-righting times. Bar graphs show X±SE values for control and treated rats, irrespective of the IDPN dose. Middle and bottom panels show the relationship between the behavioral and synaptic data. Blue dots: control rats; red dots: treated rats. Vertical dashed lines show limits defined by control and IDPN1000 rats in tail-lift angles (120 and 70 degrees, respectively, middle panels) and in air-righting times (170 and 350 ms, respectively, bottom panels). N.S.: No significant differences were found in median puncta counts among animals in the three groups defined by the two limit values in angles or times.

## 4. DISCUSSION

The vestibular system is stimulated by accelerations that largely vary in frequency and intensity. These diverse stimuli are encoded to be used in a large variety of reflex behaviors and motor control functions, as well as cognitive functions. The identification of the cellular basis of each vestibular function will contribute to our understanding of the biological roles of this system and contribute to our ability to confront vestibular deficits and diseases. In the present work, we have used a clinically relevant approach, exposure to an ototoxic compound, to cause diverse degrees of damage in the vestibular sensory epithelia, and to evaluate the relationship between the degree of damage and the functional loss in two vestibular reflexes in the rat. The results further support our conclusion that the tail-lift reflex and the air-righting reflex provide a useful quantitative measure of vestibular dysfunction (Martins-Lopes et al., 2019). Although the ototoxic treatment caused body weight loss, this side effect was transient and unlikely to affect the reflex measures.

HC counts were obtained from epithelia immunolabelled with antibodies against Caspr1 to mark HCI, calretinin to mark HCII, and Myo7a to mark all HCs. Myo7a is a well-known and widely used HC marker (Pujol et al., 2014). Caspr1 is not a marker of HCIs but can be used for this purpose because it labels the post-synaptic membrane in the inner side of the calyces (Sousa et al., 2009; Lysakowski et al., 2011; Sedó-Cabezón et al., 2015). In previous studies, we have shown that this protein can be reversibly down-regulated during chronic IDPN toxicity, an exposure model in which HC detachment from the afferent terminals and synaptic uncoupling precedes HC degeneration (Sedó-Cabezón et al., 2015; Greguske et al., 2019). However, acute or sub-acute IDPN exposure causes irreversible HC degeneration (Llorens et al., 1994; Seoane et al., 2001a,b) and the stability of the behavioral effect recorded here matches this HC loss, not a reversible loss of Caspr1. Other ototoxicity models have been reported to cause persistent calyx damage despite enduring presence of the corresponding HCI (Hirvonen et al., 2005; Sultemeier and Hoffman, 2017).In the present study, any HCI remaining in the epithelium after loss of its Caspr1+ calyx would have been counted as HCs (Myo7a+), but neither as HCI, nor as HCII. We hypothesize that these were not providing physiologically useful signals to the vestibular pathway, as illustrated by the chronic toxicity model (Sedó-Cabezón et al., 2015; Greguske et al., 2019). HCII counts were obtained using the calcium binding protein, calretinin, as marker. In the rat, the expression of calretinin has been reported to occur in 5-10 % of HCI, and from 20 % in cristae to 80 % in peripheral utricle of HCIIs (Desai et al., 2005a, b). In our specimens, we observed that levels of calretinin may vary largely, and this may have caused loss of some of these cells during segmentation of the images, causing a reduction in the estimated number of HCIIs. This may account at least in part, for the larger HCI/HCII ratios found in the present study in comparison to previous (Desai et al., 2005a,b) studies. Regarding the smaller number of all HCs compared to HCI+HCII in some crista and utricle counts from Control and IDPN400 animals, a likely explanation is the underestimation of Myo7a+ cells due to the difficulties in discerning cells in very densely packed epithelia from untreated epithelia.

Because of the known difficulty in causing vestibular HC loss in rats with the most clinically relevant ototoxins, the aminoglycoside antibiotics (Granados and Meza, 2005), we used an experimental ototoxin whose effects show many similarities with those of the aminoglycosides. These include the progression of the damage in epithelium- and zone-dependent manners. Starting with the pioneering work by Lindeman (1969), the literature contains many references, including ours on IDPN (Llorens et al., 1993; Llorens and Demêmes, 1994), stating that HCI are more sensitive to ototoxic-induced degeneration than HCII, and that the susceptibility progresses in a crista > utricle > saccule order and from the central zones to the periphery of the receptors. However, surprisingly few studies have provided quantitative assessment of these differences (Lopez et al., 1997; Nakayama et al., 1996; Hirvonen et al., 2005). In the present study, the data clearly demonstrated the higher susceptibility of HCI compared to HCII to IDPN ototoxicity, and the relative resistance of the saccule to this effect. In contrast, we found no clear differences in susceptibility to the ototoxic damage between the crista and the utricle or between the central and peripheral regions. In any case, the greater resistance of HCII and the saccule to ototoxic damage observed here likely relates to the differential susceptibility known to occur in the auditory system. Thus, in the cochlea, inner HCs are more resistant than outer HCs to aminoglycoside and cisplatin toxicity, and the susceptibility decreases progressively from the basal to the apical ends, resulting in the characteristic clinical deficits in frequency discrimination and high frequency hearing. The precise basis of these differences is not established but are believed to result from intrinsic biochemical and physiological differences among HCs (Lee et al., 2013; Fettiplace and Nam, 2019).

Increasing the doses of the toxic compound administered to the animal expectedly resulted in more extensive damage and deeper functional loss. The data allowed choosing limit values for normal responses, as defined by the behavior of control rats. In addition, comparison of histological and behavioral data allowed to define the responses corresponding to absence of vestibular input, because the IDPN1000 animals had virtually no HCs remaining. A striking outcome of the study was the low proportion of animals that showed responses intermediate between those of normal and absent vestibular input. This sharp transition between normal and totally abnormal responses was more marked in the tail-lift than in the air-righting reflexes, but in both cases undermined the power of the statistical analyses. For both reflexes, many animals showed a response identical to that of IDPN1000 rats, that is, one denoting a total absence of vestibular function, while still showing a large proportion of HCs in the vestibular epithelia. For instance, data clouds in Figure 6 reveal that tail-lift angles drop at lower doses than the doses at which HCII numbers start to decrease. One possible explanation of this observation would be that these remaining HCs are not functional. Evidence for non-functional surviving HCs has been found in an intratympanic gentamicin chinchilla model (Hirvonen et al., 2005). It is thus possible that many or all of the surviving HCs detected by immunofluorescence in this study are non-functional for sensory transduction or synaptic transmission. While this hypothesis remains to be evaluated in future studies, some data are available suggesting that surviving cells are indeed functional. First, the relationship between cell counts and reflex abnormalities were different for the two reflexes, indicating that at least some remaining function serves to one but not the other reflex. Second, although a small proportion of HCs surviving acute IDPN show stereocilia damage, most show control-like stereocilia (Llorens et al., 1993; Llorens and Demêmes, 1994; Boadas-Vaello et al., 2017), and stereociliary damage or preservation associates respectively with permanent or reversible loss of function after chronic IDPN (Sedó-Cabezón et al., 2015). Third, the normal distribution of Caspr1 label in the calyceal junctions, when these were present, also suggested a normal function, as suggested by the recovery in vestibular function associated with recovery of control-like Caspr1 label after chronic IDPN exposure (Sedó-Cabezón et al., 2015). Finally, maintenance of control-like numbers of synaptic puncta in the surviving HCII cells (Fig. 11) also supported the hypothesis that these cells may remain functional.

One alternative explanation would be that the surviving HCs remain functional but that these reflexes largely depend on a particular subset of vestibular HCs. This subset of cells would be lost at a particular toxicity level, and this would cause a fast drop in the reflex. Under this hypothesis, the present data provide initial clues to the cellular basis of the reflexes. For instance, the comparison of data in figures 5 and 6 strongly suggests that HCI have a greater role than HCII in shaping the tail-lift reflex, as the loss of this reflex seems to occur before the loss of HCII begins. Also, data in Fig. 8 suggest that HCI in the crista and utricle probably have not a major role in the air-righting reflex. Thus, only modest increases in air-righting times were recorded in animals showing a deep-to-complete loss of these cells, and times still showed dose-dependent increases after their complete loss. In relation to this point, we must also indicate that the times to right were interfered by the contact with the foam pad in the rats severely deficient in vestibular function. However, this does not invalidate the observation of nearly normal air-righting in animals bearing a deep HCI loss in the crista and utricle.

The precise role of HCI and HCII in vestibular function has not been completely established yet. However, most evidence supports the notion that the HCI/calyx units are fast adapting receptors specializing in transduction of high frequency stimuli, as needed to generate reflexes that drive fast compensatory actions required to respond to loss of balance (Eatock, 2018). The present study supports the strategy of studying the impact of partial vestibular lesions on behavior to establish hypotheses on the ultimate physiological roles of hair cell types and vestibular sub-systems. Future studies using this approach may benefit from the increasing knowledge of cell and afferent subtypes subdivided by zones as defined by the expression of unique proteins (McInturff et al., 2018; Hoffman et al., 2018), as well as from the use of more complex approaches, such as genetic manipulations (Ono et al., 2020), and the integration of afferent recording (Hirvonen et al., 2005).

Another aspect in need of future research is the extrapolation to other species, most notably the mouse. In view of the known combinations of common features and species-specific differences in vestibular histology and physiology (Boyle et al., 1992; Curthoys et al., 2017; Desai et al., 2005a,b; Eatock, 2018; Goldberg, 2000), it would be premature to assume that the relationship between reflex loss and HC loss will be in the mouse like in the rat. For instance, ventral curling during tail-lift has been reported in many vestibular-deficient mice, but our observations after vestibular toxicity in rats (Llorens et al., 1993; Llorens and Rodríguez-Farré., 1997; Martins-Lopes et al., 2019; this study) and mice (Soler-Martin et al., 2007; Saldaña-Ruíz et al., 2013; Boadas-Vaello et al., 2017; Greguske et al., 2019) suggest that the abnormal tail-lift response more easily and robustly develops in the former than the latter species.

## 5. CONCLUSION

The present study compared the loss of vestibular HCs with the loss of anti-gravity reflex responses in rats following graded ototoxicity. While both the tail-lift reflex and the air-righting reflex were affected in a dose-dependent manner, the former was slightly more sensitive than the latter to the toxic damage, suggesting that these two reflexes depend on non-identical populations of HCs. Loss of the tail-lift reflex better associated with loss of HCI, suggesting that this reflex may predominantly depend on HCI function. Additional studies are needed to corroborate this hypothesis, including an evaluation of the functional competence of the surviving cells in the partially damaged vestibular epithelia.

## Supporting information

Legends for Supplementary Figures

## Acknowledgements

This study was supported by grants RTI2018-096452-B-I00 (Ministerio de Ciencia, Innovación y Universidades, Agencia Estatal de Investigación, Fondo Europeo de Desarrollo Regional, MCIU/AEI/FEDER, UE), and 2017 SGR 621 (Agència de Gestió d’Ajuts Universitaris i de Recerca, Generalitat de Catalunya). A.B.G is a Serra-Húnter fellow. The confocal microscopy studies were performed at the Scientific and Technological Centers of the University of Barcelona (CCiT-UB). We thank Dr. Benjamin Torrejon for technical assistance, and former students of the laboratory, Anna Bellmunt and Vanessa Martins-Lopes, for their contributions to the study. We also thank Drs. Ruth Anne Eatock and Kazuya Ono for their critical review of the manuscript.

## REFERENCES

Baird RA, Desmadryl G, Fernández C, Goldberg JM (1988) The vestibular nerve of the chinchilla. II. Relation between afferent response properties and peripheral innervation patterns in the semicircular canals. J Neurophysiol 60: 182–203.

Beraneck M, Bojados M, Le Séac’h A, Jamon M, Vidal PP (2012) Ontogeny of mouse vestibuloocular reflex following genetic or environmental alteration of gravity sensing. PLoS One 7: e40414. doi: 10.1371/journal.pone.0040414.

Boadas-Vaello P, Sedó-Cabezón L, Verdú E, Llorens J (2017) Strain and Sex Differences in the Vestibular and Systemic Toxicity of 3,3’-Iminodipropionitrile in Mice. Toxicol Sci 156: 109–122.

Boyle R, Goldberg JM, Highstein SM (1992) Inputs from regularly and irregularly discharging vestibular nerve afferents to secondary neurons in squirrel monkey vestibular nuclei. III. Correlation with vestibulospinal and vestibuloocular output pathways. J Neurophysiol 68: 471–484.

Cassel R, Bordiga P, Carcaud J, Simon F, Beraneck M, Le Gall A, Benoit A, Bouet V, Philoxene B, Besnard S, Watabe I, Pericat D, Hautefort C, Assie A, Tonetto A, Dyhrfjeld-Johnsen J, Llorens J, Tighilet B, Chabbert C (2019) Morphological and functional correlates of vestibular synaptic deafferentation and repair in a mouse model of acute-onset vertigo. Dis Model Mech 12(7). pii: dmm039115. doi: 10.1242/dmm.039115.

Corneil BD, Camp AJ (2018) Animal models of vestibular evoked myogenic potentials: The past, present, and future. Front Neurol 9: 489. doi: 10.3389/fneur.2018.00489.

Crofton KM, Knight T (1991) Auditory deficits and motor dysfunction following iminodipropionitrile administration in the rat. Neurotoxicol Teratol. 13: 575–581.

Crofton KM, Janssen R, Prazma J, Pulver S, Barone Jr. S (1994) The ototoxicity of 3,3’-iminodipropionitrile: Functional and morphological evidence of cochlear damage. Hear Res 80: 129–140.

Curthoys IS, Vulovic V, Manzari L (2012) Ocular vestibular-evoked myogenic potential (oVEMP) to test utricular function: neural and oculomotor evidence. Acta Otorhinolaryngol Ital. 32: 41–45.

Curthoys IS, Grant JW, Burgess AM, Pastras CJ, Brown DJ, Manzari L (2018) Otolithic receptor mechanisms for vestibular-evoked myogenic potentials: A review. Front Neurol 9: 366. doi: 10.3389/fneur.2018.00366.

Curthoys IS, MacDougall HG, Vidal P-P, de Waele C (2017) Sustained and transient vestibular systems: A physiological basis for interpreting vestibular function. Front Neurol 8: 117. doi: 10.3389/fneur.2017.00117.

De Jeu M, De Zeeuw CI (2012) Video-oculography in mice. J Vis Exp 65: e3971. doi: 10.3791/3971.

Dechesne CJ, Winsky L, Kim HN, Goping G, Vu TD, Wenthold RJ, Jacobowitz DM (1991) Identification and ultrastructural localization of a calretinin-like calcium-binding protein (protein 10) in the guinea pig and rat inner ear. Brain Res 560, 139–148

Desai SS, Zeh C, Lysakowski A. (2005a) Comparative morphology of rodent vestibular periphery. I. Saccular and utricular maculae. J Neurophysiol 93: 251–266.

Desai SS, Ali H, Lysakowski A (2005b) Comparative morphology of rodent vestibular periphery. II. Cristae ampullares. J Neurophysiol 93: 267–280.

Desmadryl G, Dechesne CJ (1992) Calretinin immunoreactivity in chinchilla and guinea pig vestibular end organs characterizes the calyx unit subpopulation. Exp Brain Res 89, 105–108.

Eatock RA (2018) Specializations for fast signaling in the amniote vestibular inner ear. Integr Comp Biol 58: 341–350.

Eatock RA, Songer JE (2011) Vestibular hair cells and afferents: two channels for head motion signals. Annu Rev Neurosci 34: 501–534.

Fettiplace R, Nam JH (2019) Tonotopy in calcium homeostasis and vulnerability of cochlear hair cells. Hear Res 376: 11–21.

Goldberg JM (2000) Afferent diversity and the organization of central vestibular pathways. Exp Brain Res 130: 277–297.

Granados O, Meza G (2005) Streptidine, a metabolic derivative produced after administration of streptomycin in vivo, is vestibulotoxic in rats. Histol Histopathol 20: 357–364.

Greguske EA, Carreres-Pons M, Cutillas B, Boadas-Vaello P, Llorens J (2019) Calyx junction dismantlement and synaptic uncoupling precede hair cell extrusion in the vestibular sensory epithelium during sub-chronic 3,3’-iminodipropionitrile ototoxicity in the mouse. Arch Toxicol 93: 417–434.

Hasson T, Gillespie PG, Garcia JA, MacDonald RB, Zhao Y, Yee AG, Mooseker MS, Corey DP (1997) Unconventional myosins in inner-ear sensory epithelia. J Cell Biol 137, 1287–1307.

Halmagyi GM, Chen L, MacDougall HG, Weber KP, McGarvie LA, Curthoys IS (2017) The video head impulse test. Front Neurol 8: 258. doi: 10.3389/fneur.2017.00258.

Hirvonen TP, Minor LB, Hullar TE, Carey JP (2005) Effects of intratympanic gentamicin on vestibular afferents and hair cells in the chinchilla. J Neurophysiol 93, 643–655.

Hoffman LF, Choy KR, Sultemeier DR, Simmons DD (2018) Oncomodulin expression reveals new insights into the cellular organization of the murine utricle striola. J Assoc Res Otolaryngol 19, 33–51.

Hunt M.A., Miller S.W., Nielson H.C., Horn K.M. (1987) Intratympanic injections of sodium arsanilate (atoxil) solution results in postural changes consistent with changes described for labyrinthectomized rats. Behav Neurosci 101, 427–428.

Imai T, Takimoto Y, Takeda N, Uno A, Inohara H, Shimada S (2016) High-speed video-oculography for measuring three-dimensional rotation vectors of eye movements in mice. PLoS One 11: e0152307. doi: 10.1371/journal.pone.0152307.

Lee JH, Park C, Kim SJ, Kim HJ, Oh GS, Shen AH, So HS, Park R (2013) Different uptake of gentamicin through TRPV1 and TRPV4 channels determines cochlear hair cell vulnerability. Exp Mol Med 45: e12. doi: 10.1038/emm.2013.25.

Lindeman H.H. (1969) Regional differences in sensitivity of the vestibular sensory epithelia to ototoxic antibiotics. Acta Otolaryngol 67: 177–189.

Llorens J, Demêmes D (1994) Hair cell degeneration resulting from 3,3’-iminodipropionitrile toxicity in the rat vestibular epithelia. Hear Res 76: 78–86.

Llorens J, Demêmes D, Sans A (1993) The behavioral syndrome caused by 3,3’-iminodipropionitrile and related nitriles in the rat is associated with degeneration of the vestibular sensory hair cells. Toxicol Appl Pharmacol 123:199–210.

Llorens J, Rodríguez-Farré E (1997) Comparison of behavioral, vestibular, and axonal effects of subchronic IDPN in the rat. Neurotoxicol Teratol 19: 117–127.

Lysakowski A, Gaboyard-Niay S, Calin-Jageman I, Chatlani S, Price SD, Eatock RA (2011) Molecular microdomains in a sensory terminal, the vestibular calyx ending. J Neurosci 31: 10101–10114.

Lopez I, Honrubia V, Lee S.C., Schoeman G, Beykirch K (1997) Quantification of the process of hair cell loss and recovery in the chinchilla crista ampullaris after gentamicin treatment. Int J Dev Neurosci 15: 447–461.

Maroto AF, Greguske EA, Deulofeu M, Boadas-Vaello P, Llorens J (2021) Behavioral assessment of vestibular dysfunction in rats. In: Experimental Neurotoxicology Methods, Neuromethods, vol 172, Llorens J and Barenys M Editors. Springer US, pp xxx–xxx

Martins-Lopes V, Bellmunt A, Greguske EA, Maroto AF, Boadas-Vaello P, Llorens J (2019) Quantitative assessment of anti-gravity reflexes to evaluate vestibular dysfunction in rats. J Assoc Res Otolaryngol 20: 553–563.

McInturff S, Burns JC, Kelley MW (2018) Characterization of spatial and temporal development of Type I and Type II hair cells in the mouse utricle using new cell-type-specific markers. Biol Open 7: bio038083. doi: 10.1242/bio.038083.

Nakayama M, Riggs LC, Matz GJ (1996) Quantitative study of vestibulotoxicity induced by gentamicin or cisplatin in the guinea pig. Laryngoscope. 106: 162–167.

Ono K, Keller J, López Ramírez O, González Garrido A, Zobeiri OA, Chang HHV, Vijayakumar S, Ayiotis A, Duester G, Della Santina CD, Jones SM, Cullen KE, Eatock RA, Wu DK. (2020) Retinoic acid degradation shapes zonal development of vestibular organs and sensitivity to transient linear accelerations. Nat Commun 11: 63. doi: 10.1038/s41467-019-13710-4.

Pellis SM, Pellis VC, Morrissey TK, Teitelbaum P (1989) Visual modulation of vestibularly-triggered air-righting in the rat. Behav Brain Res 35: 23–26.

Pellis SM, Pellis VC, Teitelbaum P (1991) Labyrinthine and other supraspinal inhibitory controls over head-and-body ventroflexion. Behav Brain Res. 46: 99–102.

Pujol R, Pickett SB, Nguyen TB, Stone JS (2014) Large basolateral processes on type II hair cells are novel processing units in mammalian vestibular organs. J Comp Neurol 522:3141–3159.

Saldaña-Ruíz S, Boadas-Vaello P, Sedó-Cabezón L, Llorens J (2013) Reduced systemic toxicity and preserved vestibular toxicity following co-treatment with nitriles and CYP2E1 inhibitors: a mouse model for hair cell loss. J Assoc Res Otolaryngol 14: 661–671.

Sedó-Cabezón L, Jedynak P, Boadas-Vaello P, Llorens J (2015) Transient alteration of the vestibular calyceal junction and synapse in response to chronic ototoxic insult in rats. Dis Model Mech 8: 1323–1337.

Seoane A, Demêmes D, Llorens J (2001a). Relationship between insult intensity and mode of hair cell loss in the vestibular system of rats exposed to 3,3’-iminodipropionitrile. J Comp Neurol 439: 385–399.

Seoane A, Demêmes D, Llorens J (2001b). Pathology of the rat vestibular sensory epithelia during subchronic 3,3’-iminodipropionitrile exposure: hair cells may not be the primary target of toxicity. Acta Neuropathol 102: 339–348.

Soler-Martín C, Diez-Padrisa N, Boadas-Vaello P, Llorens J (2007) Behavioral disturbances and hair cell loss in the inner ear following nitrile exposure in mice, guinea pigs, and frogs. Toxicol Sci 96: 123–132.

Sousa AD, Andrade LR, Salles FT, Pillai AM, Buttermore ED, Bhat MA, Kachar B (2009) The septate junction protein caspr is required for structural support and retention of KCNQ4 at calyceal synapses of vestibular hair cells. J Neurosci 29: 3103–3108.

Sultemeier DR, Hoffman LF (2017) Partial aminoglycoside lesions in vestibular epithelia reveal broad sensory dysfunction associated with modest hair cell loss and afferent calyx retraction. Front Cell Neurosci 11: 331. doi: 10.3389/fncel.2017.00331.

Wilkerson BA, Artoni F, Lea C, Ritchie K, Ray CA, Bermingham-McDonogh O (2018) Effects of 3,3’-iminodipropionitrile on hair cell numbers in cristae of CBA/CaJ and C57BL/6J Mice. J Assoc Res Otolaryngol 19: 483–491.

