## Supplementary Figures for "Relationship between vestibular hair cell loss and deficits in two anti-gravity reflexes in the rat"

Supplementary Figure S1

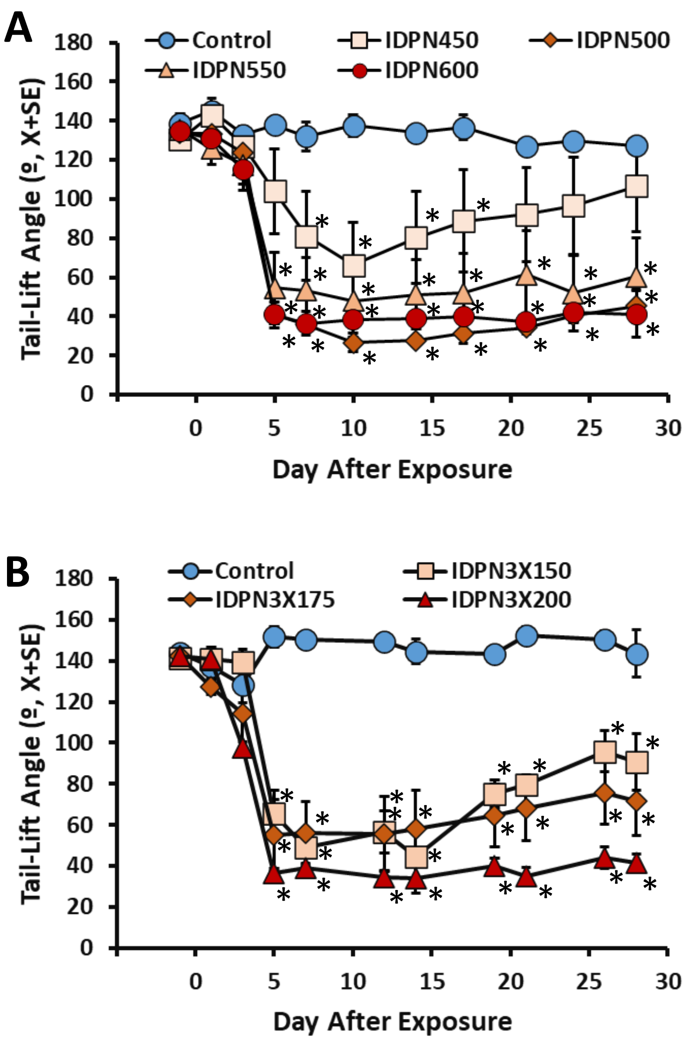

Supplementary Figure S2

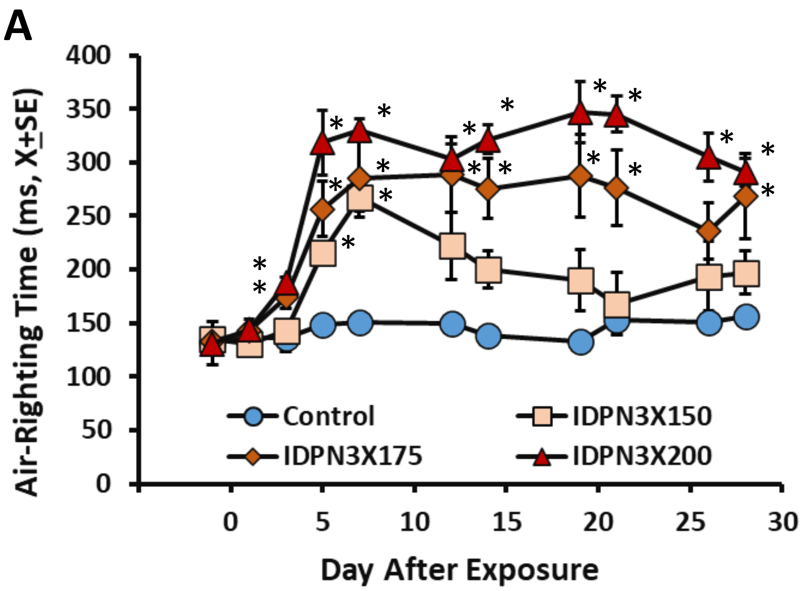

### Supplementary Figure S3

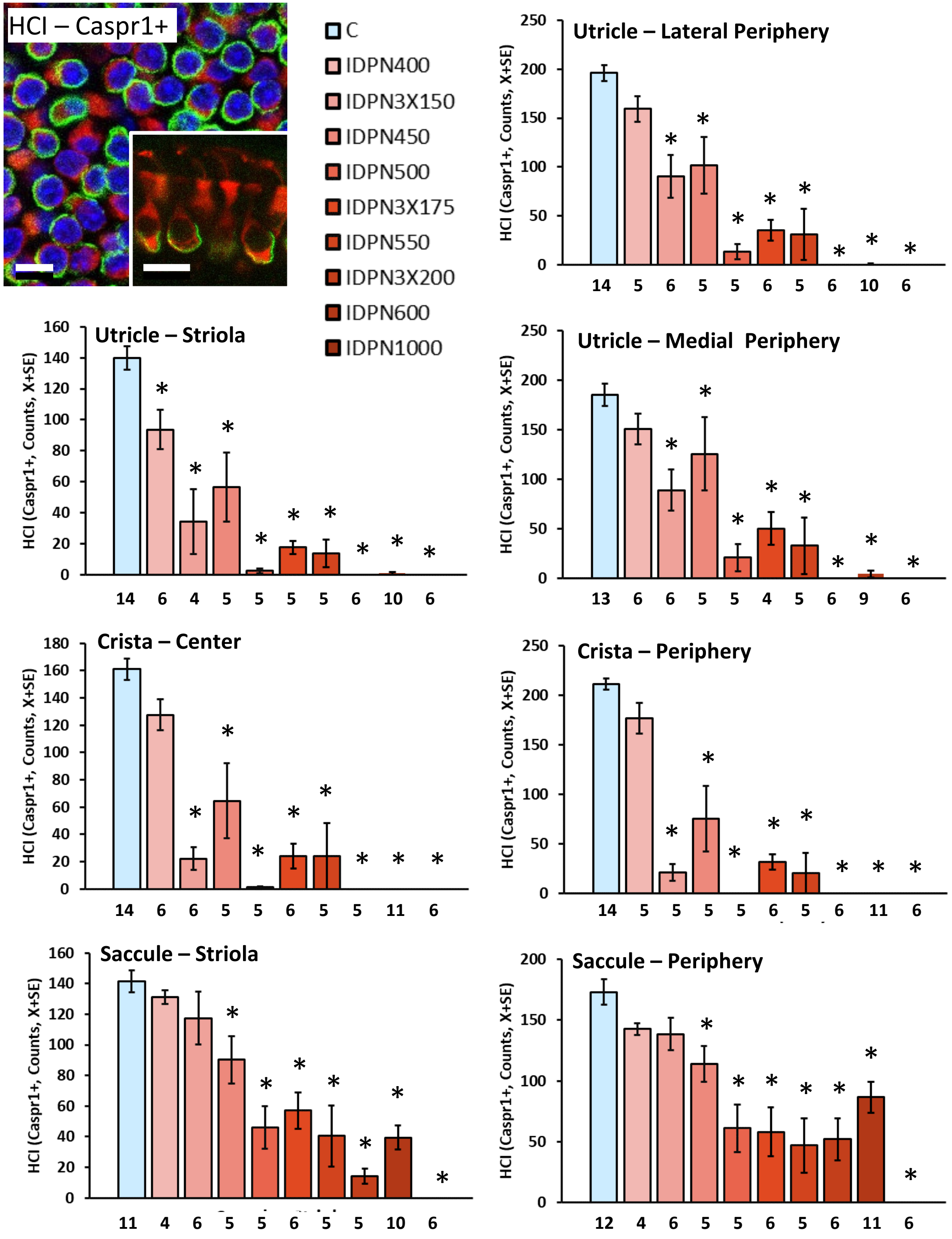

Supplementary Figure S4

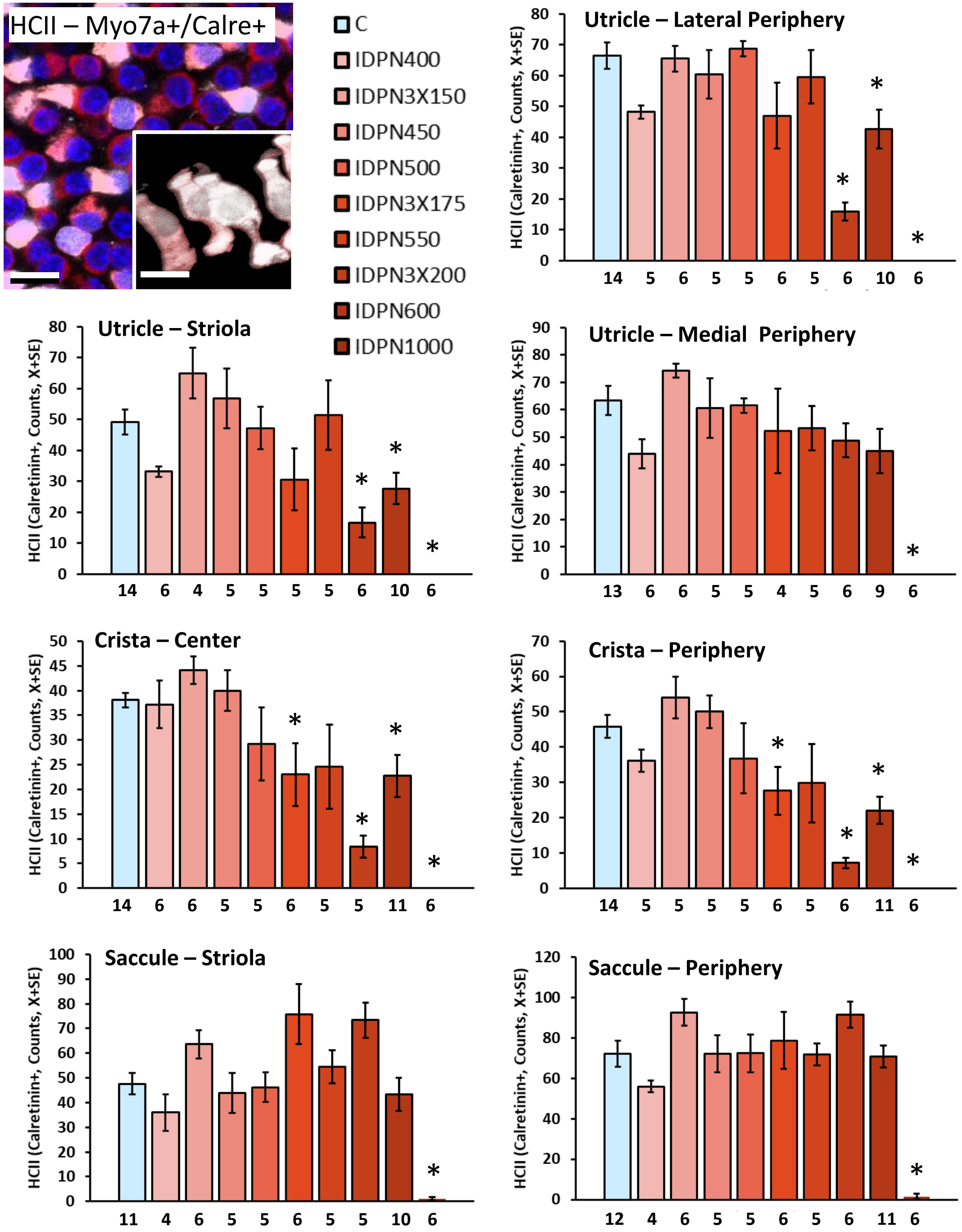

### Supplementary Figure S5

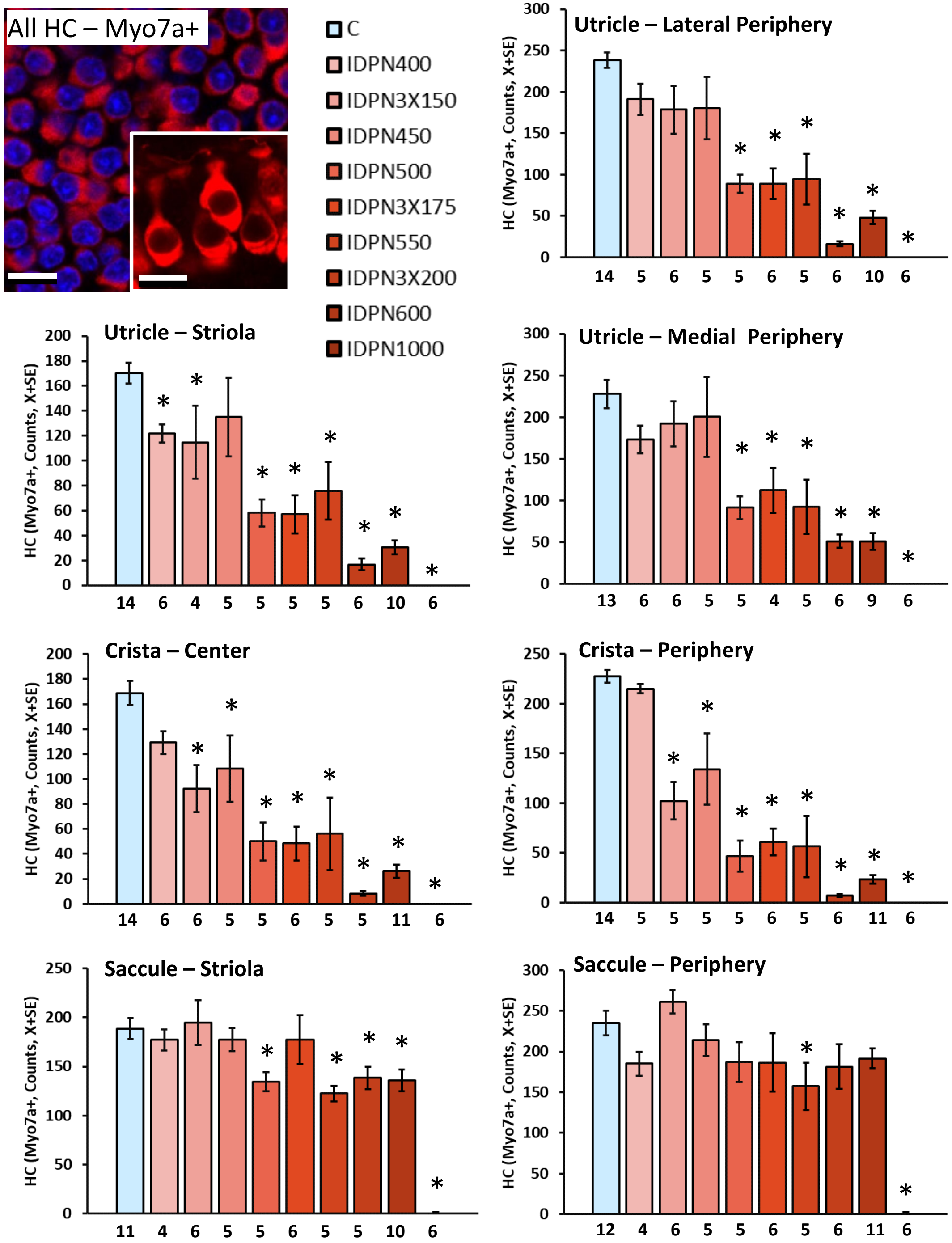

### Supplementary Figure S6

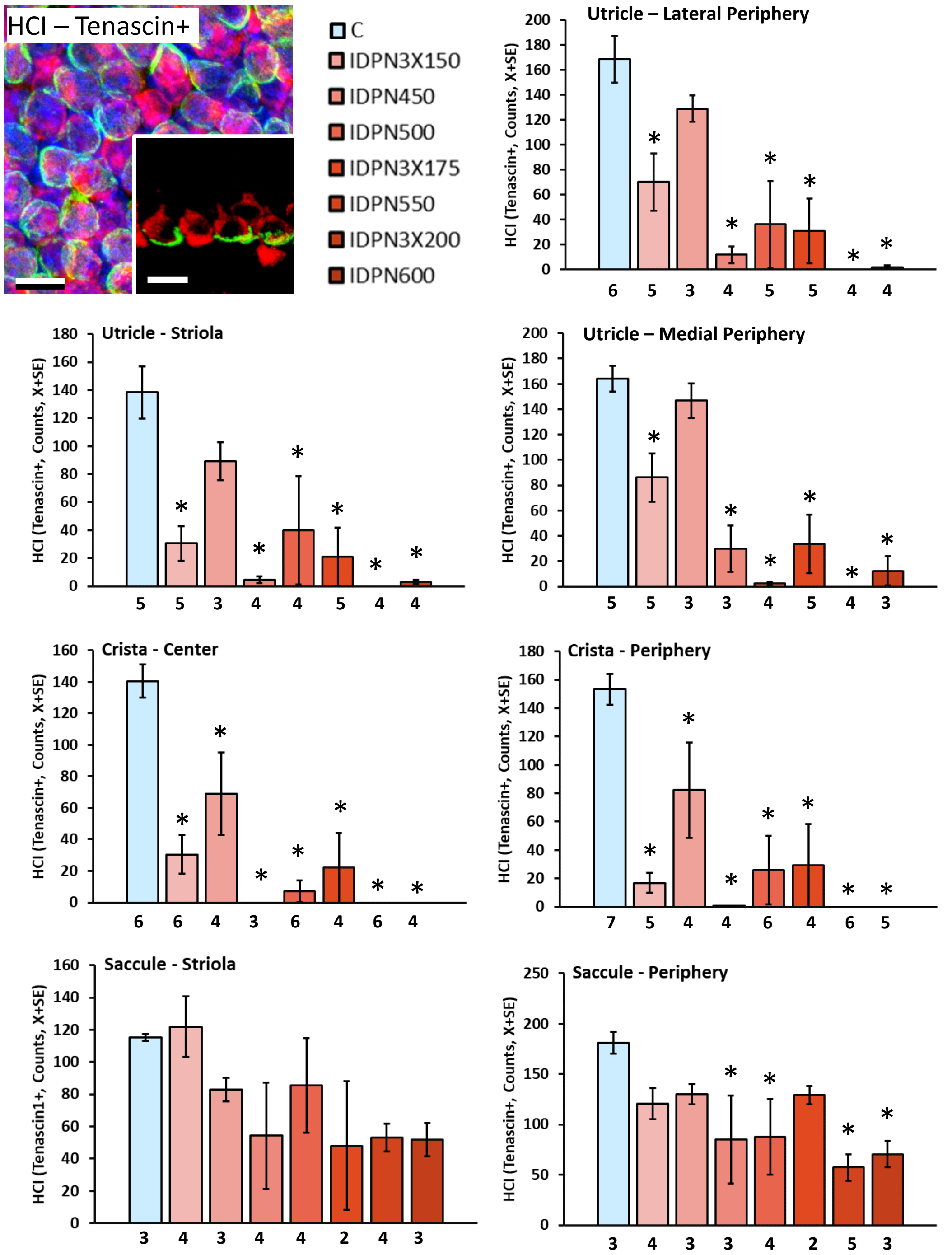

Supplementary Figure S7

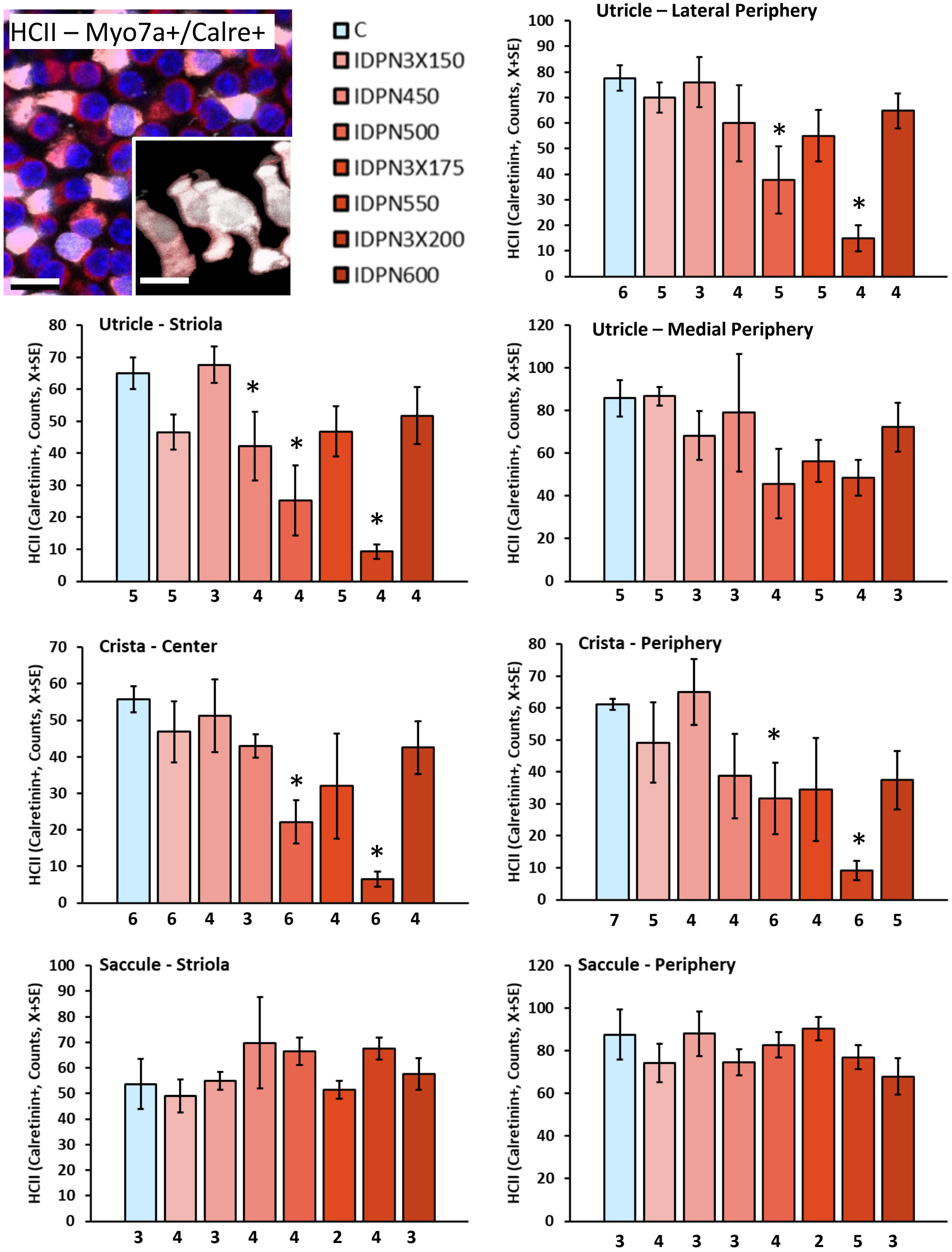

Supplementary Figure S8

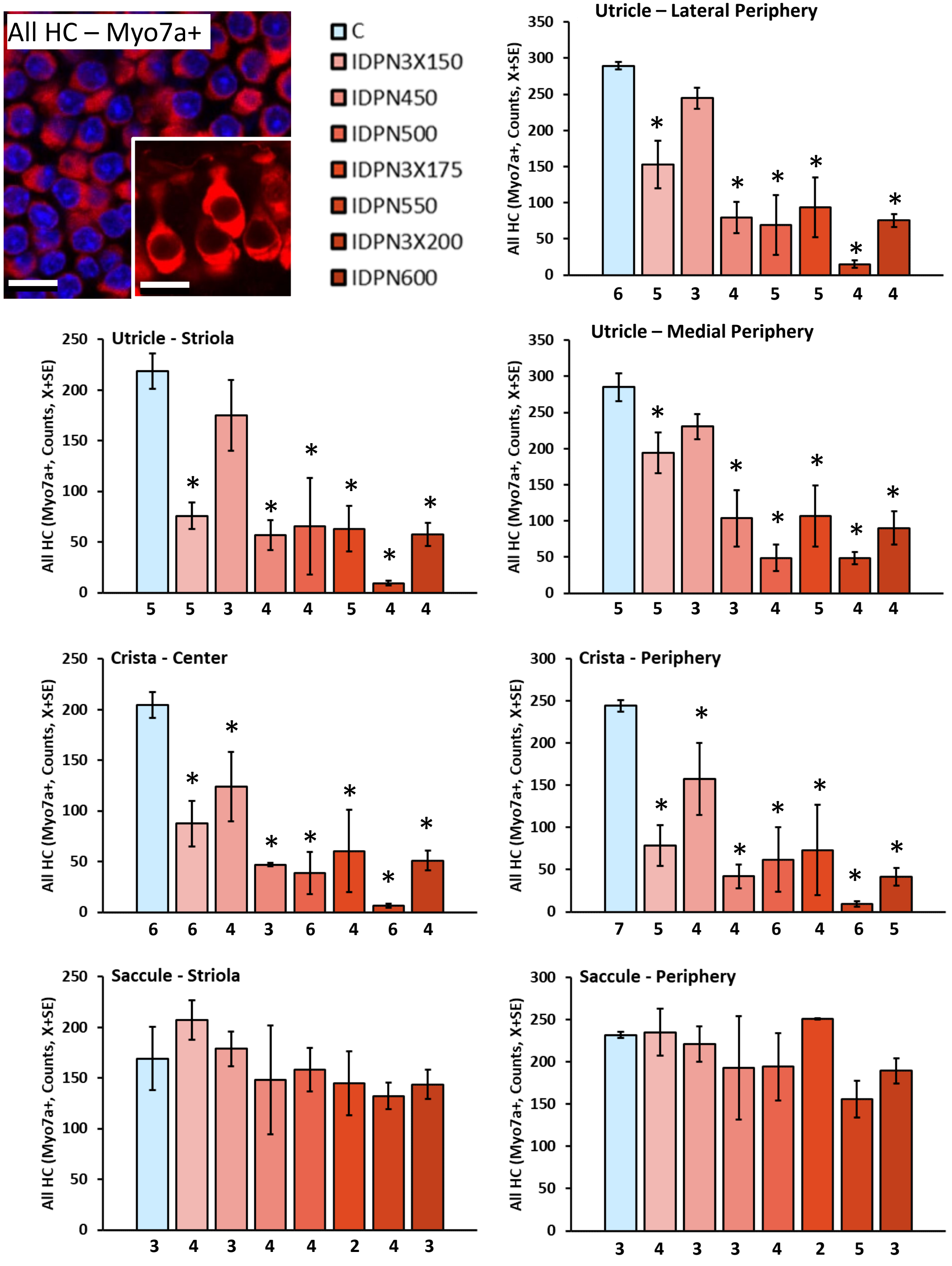

Supplementary Figure S9

**HCI (Tenascin+)**
**Tail-Lift Angle**

(n= 27-41)

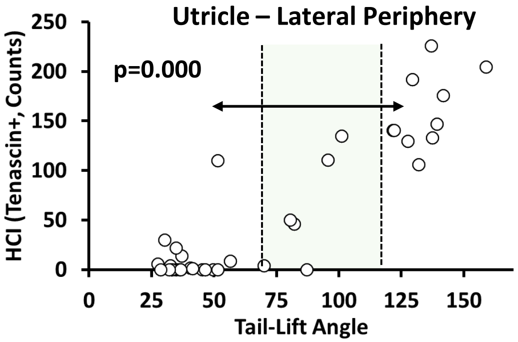


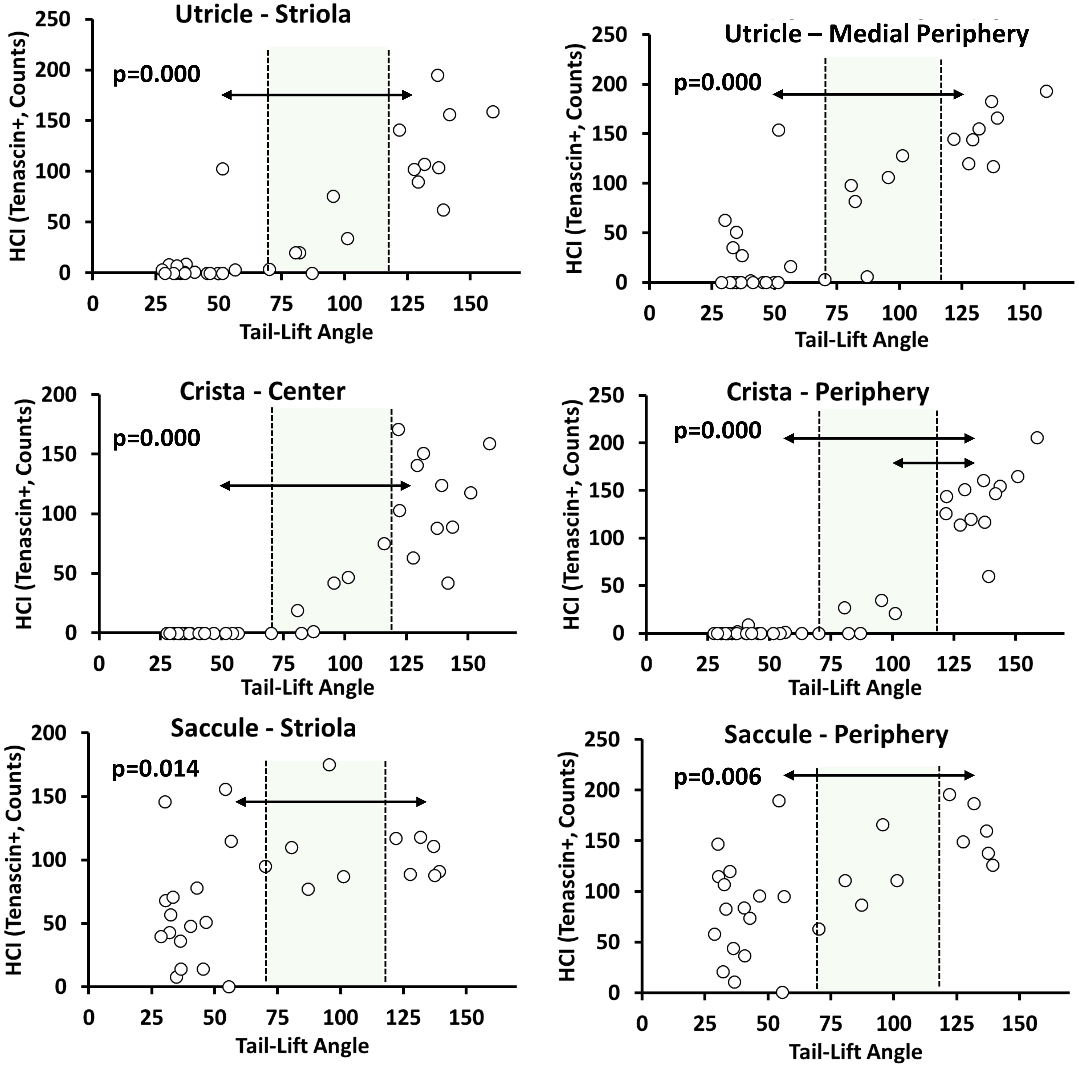

Supplementary Figure S10

**HCII (Myo7a+/Calre+)**
**Tail-Lift Angle**

(n= 27-41)

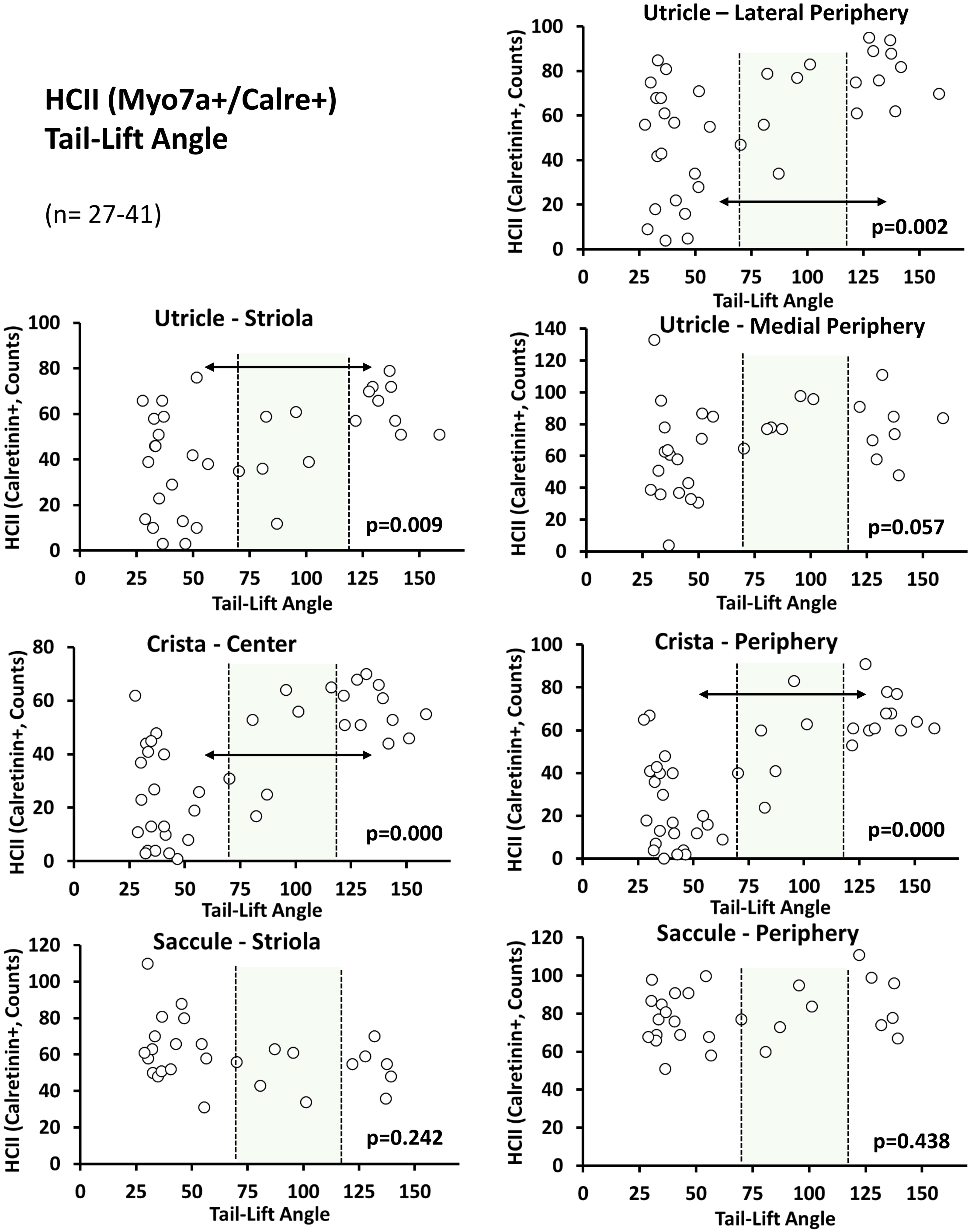

Supplementary Figure S11

**All HC (Myo7a+)**
**Tail-Lift Angle**

(n= 27-41)

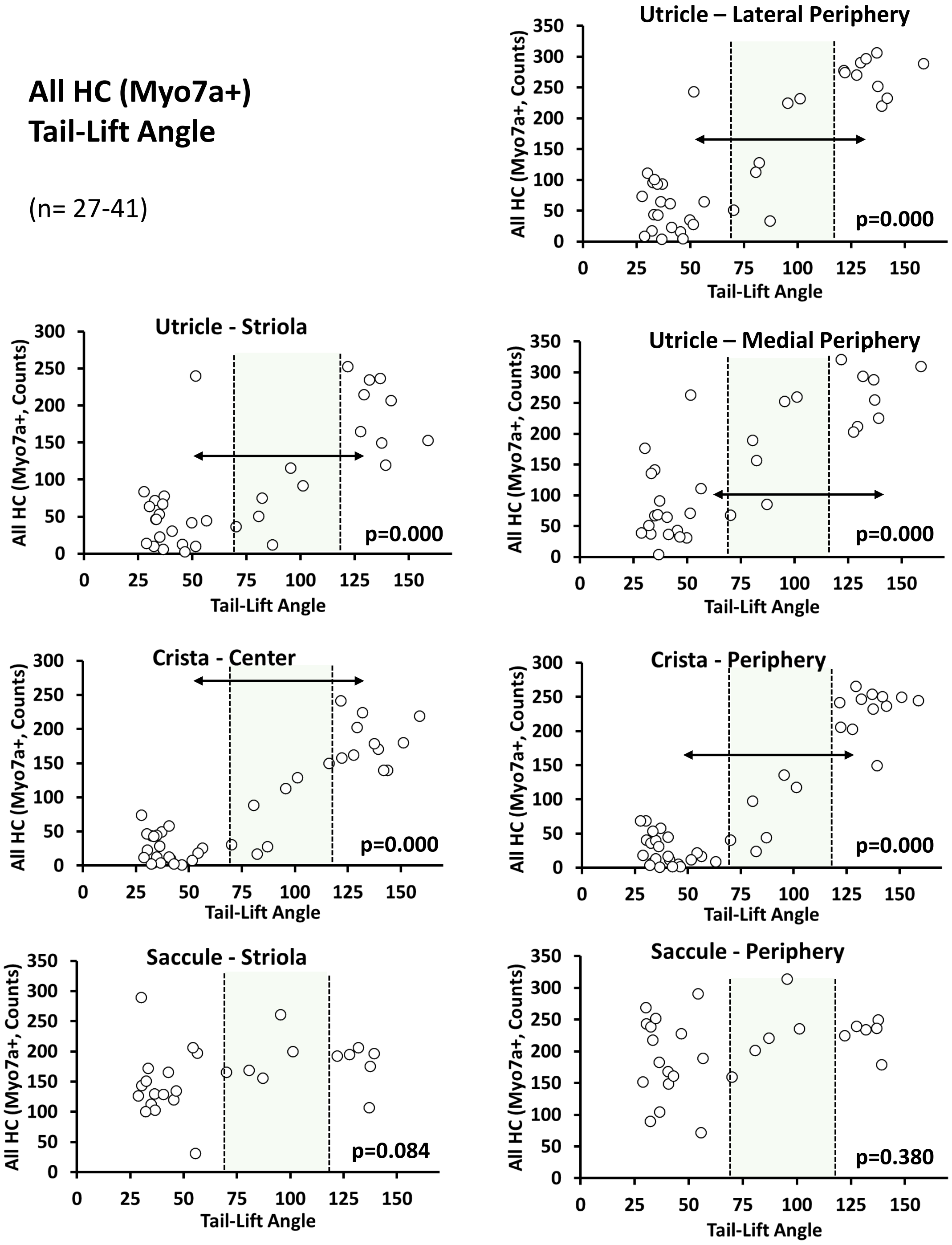

Supplementary
Figure S12

**HCI (Tenascin+)
Air-Righting Time**

(n= 13-20)

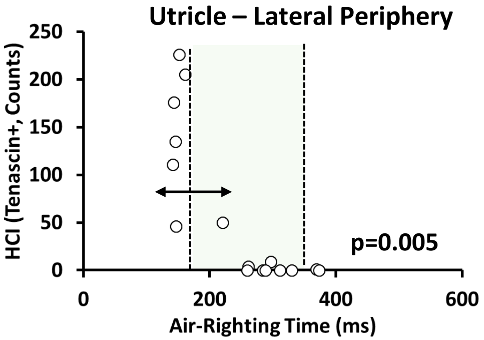


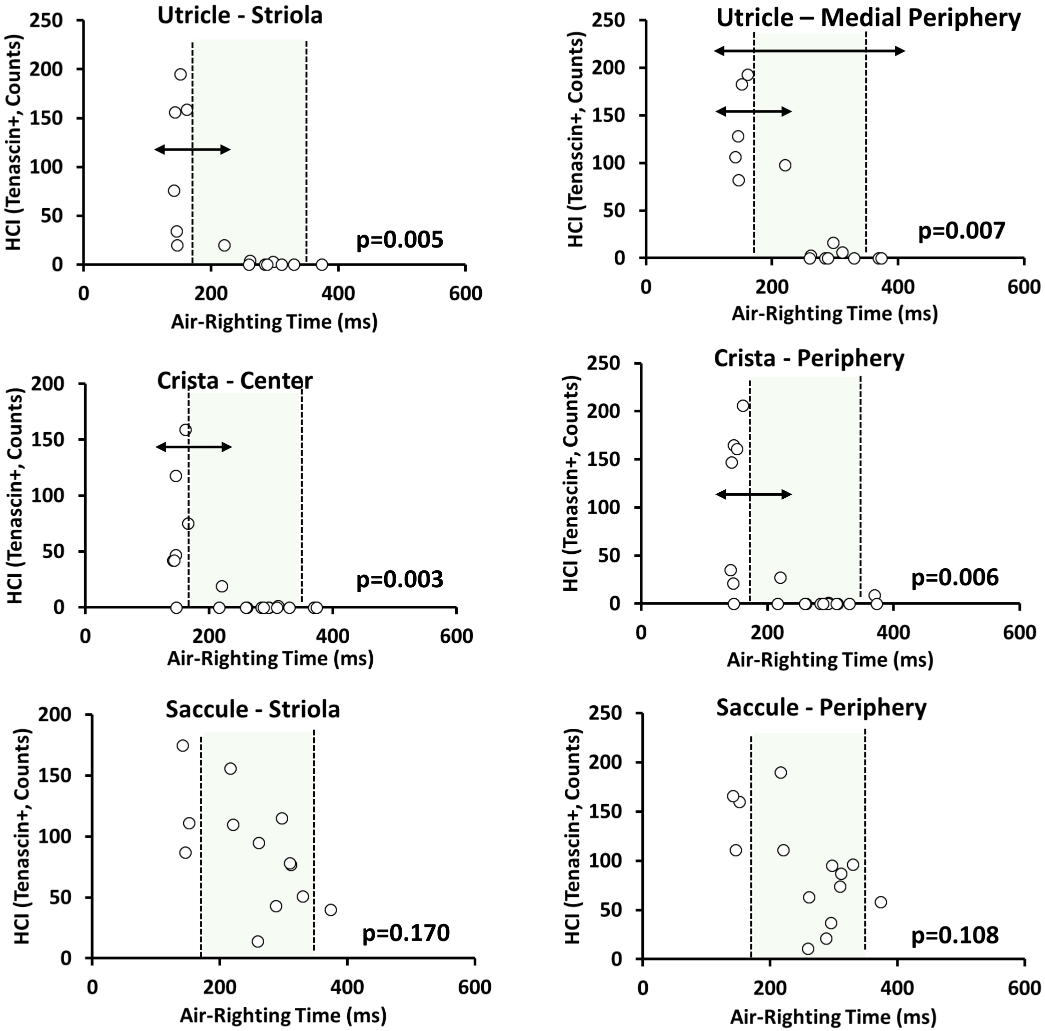

Supplementary Figure S13

**HCII (Myo7a+/Calre+) Air-Righting Time**

(n= 13-20)

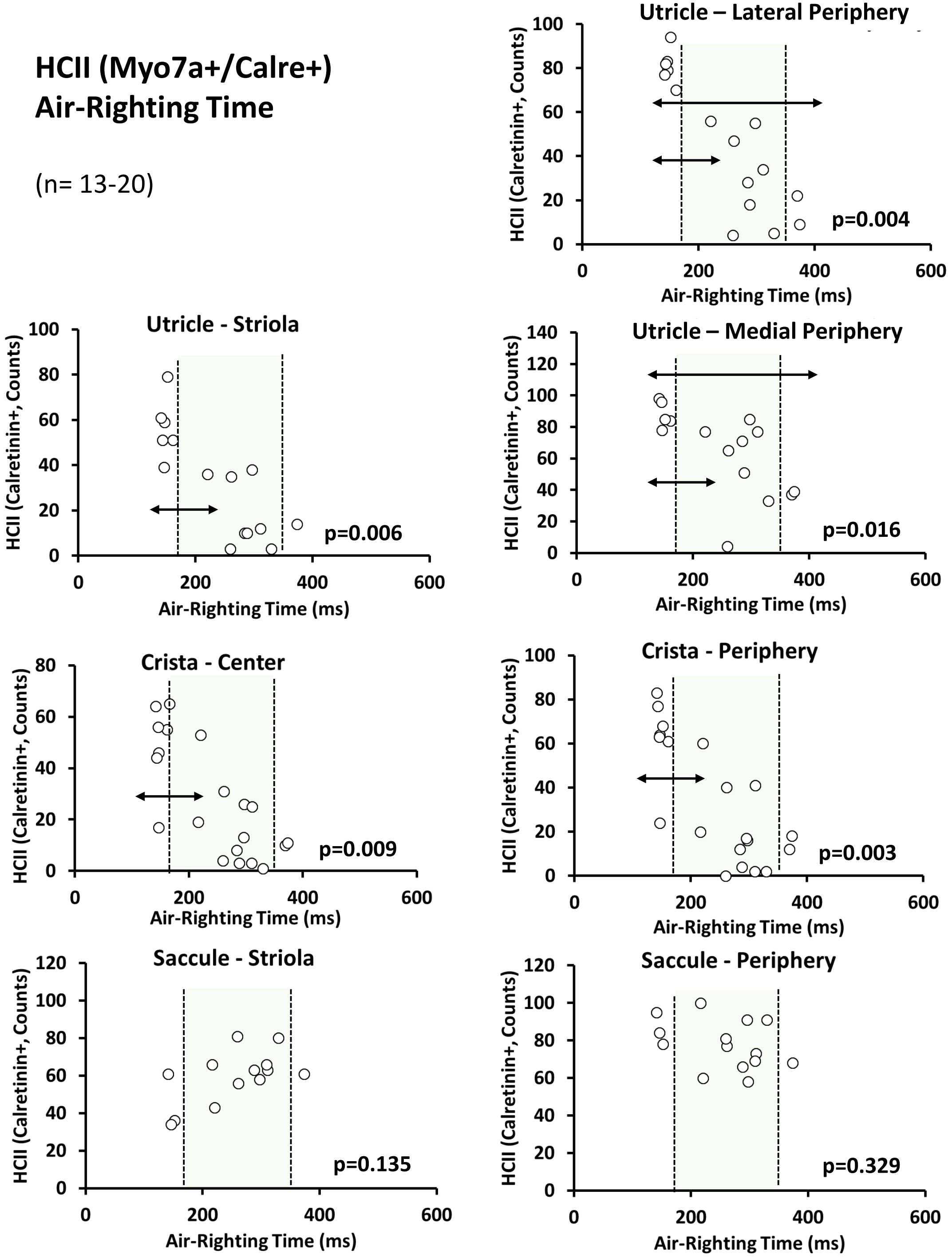

Supplementary Figure S14

**All HC (Myo7a+)**
**Air-Righting Time**

(n= 13-20)

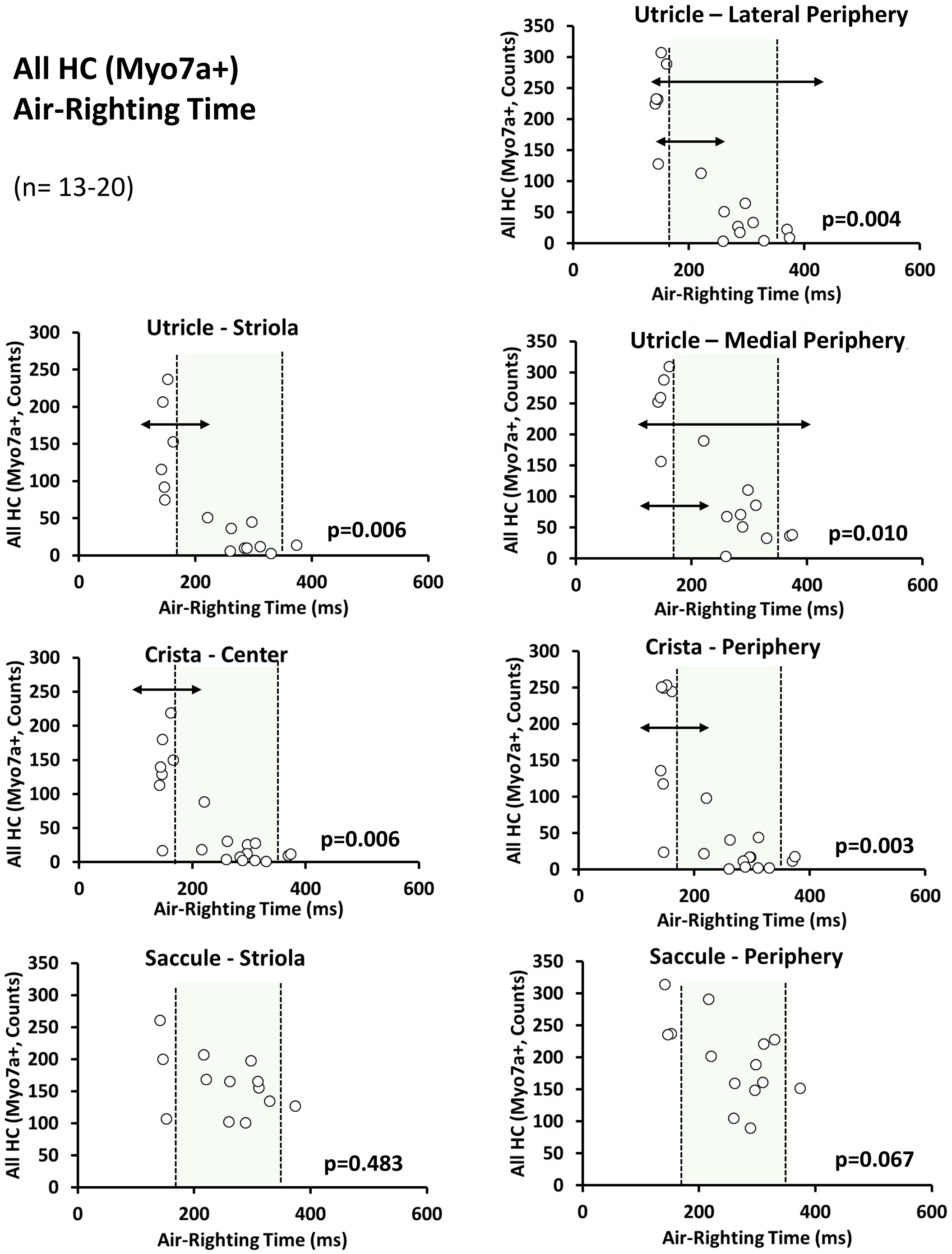
