## Supplementary material for "Relationship between vestibular hair cell loss and deficits in two anti-gravity reflexes in the rat": Legends for Supplementary Figures

LEGENDS FOR SUPPLEMENTARY MOVIES AND FIGURES

Supplementary Movies. A. Tail-lift reflex test, control rat. B. Tail-lift reflex, vestibular-deficient rat. C. Air-righting reflex test, control rat. D. Air-righting reflex test, vestibular-deficient rat.

Figure S1. Effects of vestibular toxicity on the tail-lift reflex. Data are X±SE minimum nose-neck-base of the tail angles displayed by the rats when lifted by the tail and lowered back. (A) Time course of the effect of acute IDPN, experiment 2 (0 - 600 mg/kg, n=5/group). (B) Time course of the effect of subacute IDPN, experiment 3 (0 - 200 mg/kg·day, for 3 consecutive days, n=6/group). *: p<0.05, significantly different from control group, Duncan's test after significant ANOVA and repeated-measures MANOVA analyses. Repeated-measures MANOVA analysis, with day as the within-subject factor, resulted in significant Day and Treatment effects in the tail-lift angle values recorded in experiment 2 (Panel A; Day: Wilks’ lambda = 0.070, F[10, 11] = 14.60, p = 0.000; Treatment: F[4, 20] = 8.89, p = 0.000) and experiment 3 (Panel B; Day: Wilks’ lambda = 0.088, F[10, 8] = 8.25, p = 0.003; Treatment: F[3, 17] = 17.08, p = 0.000). Day by day ANOVA analyses indicated that significant effects on tail-lift angles occurred at day 5 post-exposure and all later days in both experiments (all Fs [4, 20] > 6.43, p’s < 0.002; and all Fs [3, 17] > 8.03, p’s < 0.002). Total doses of 500 mg/kg of IDPN or more caused a maximal drop in angle by day 5 after exposure with no subsequent recovery. In animals treated with an acute dose of 450 mg/kg the maximal effect was observed by two weeks after administration. This was followed by significant recovery and the mean group value was not significantly different from control mean four weeks after administration. Animals dosed with IDPN3x150 displayed some apparent recovery in tail-lift angle values, but the mean group values did not recover to control values.

Figure S2. Effects of vestibular toxicity on the air-righting reflex. Data are X±SE air-righting times displayed by the rats when dropped in supine position from approximately 40 cm above a foam cushion. The graph shows the time course of the effect of subacute IDPN, experiment 3 (0 - 200 mg/kg·day, for 3 consecutive days, n=6/group). *: p<0.05, significantly different from control group, Duncan's test after significant ANOVA and repeated-measures MANOVA analyses. Repeated-measures MANOVA analysis, with day as the within-subject factor, revealed significant Day (Wilks’ lambda = 0.081, F[10, 8] = 9.063, p = 0.002) and Treatment (F[3, 17] = 12.18, p = 0.000) effects. Day by day ANOVA analyses indicated that significant differences among groups were present at all days between day 3 after exposure and the end of the experiment (all Fs [3, 17] > 4.58, p’s < 0.016). Maximal effects were recorded by day 7 after exposure. This was followed by significant recovery in the IDPN3X150 animals, but not in the IDPN3X175 and the IDPN3X200 animals.

Figure S3. Differences in the loss of type I hair cells (HCI) after exposure to the ototoxic compound, IDPN, as a function of the dose (400 to 1000 mg/kg, see legend), zone (central/striola vs periphery) and end-organ (utricle, crista and saccule). The images in the upper-left corner show the use of the Myo7a (red) and Caspr1 (green) labels to identify HCI. Scale bars = 10 μm. Bar graphs show HCI counts (X±SE) according to end-organ, region within it and IDPN dose. Numbers below the bars indicate numbers of animals. *: p<0.05, significantly different from control group, Duncan's test after significant ANOVA. Data in this figure are also shown together with data in Supplementary figures S4 and S5 as main Fig. 4.

Figure S4. Differences in the loss of type II hair cells (HCII) after exposure to the ototoxic compound, IDPN, as a function of the dose (400 to 1000 mg/kg, see legend), zone (central/striola vs periphery) and end-organ (utricle, crista and saccule). The images in the upper-left corner show the use of the Myo7a (red) and calretinin (white) labels to identify HCII. Scale bars = 10 μm. Bar graphs show HCII counts (X±SE) according to end-organ, region within it and IDPN dose. Numbers below the bars indicate numbers of animals. *: p<0.05, significantly different from control group, Duncan's test after significant ANOVA. Data in this figure are also shown together with data in Supplementary figures S3 and S5 as main Fig. 4.

Figure S5. Differences in the loss of all hair cells (HC) after exposure to the ototoxic compound, IDPN, as a function of the dose (450 to 600 mg/kg, see legend), zone (central/striola vs periphery) and end-organ (utricle, crista and saccule). The images in the upper-left corner show the use of the Myo7a (red) label to identify HCs. Scale bars = 10 μm. Bar graphs show HC counts (X±SE) according to end-organ, region within it and IDPN dose. Numbers below the bars indicate numbers of animals. *: p<0.05, significantly different from control group, Duncan's test after significant ANOVA. Mean data in this figure are also shown together with data in Supplementary figures S3 and S4 as main Fig. 4.

Figure S6. Differences in the loss of type I hair cells (HCI) after exposure to the ototoxic compound, IDPN, as a function of the dose (450 to 600 mg/kg, see legend), zone (central/striola vs periphery) and end-organ (utricle, crista and saccule). The images in the upper-left corner show the use of the Myo7a (red) and Tenascin (green) labels to identify HCI. Scale bars = 10 μm. Bar graphs show HCI counts (X±SE) according to end-organ, region within it and IDPN dose. Numbers below the bars indicate numbers of animals. *: p<0.05, significantly different from control group, Duncan's test after significant ANOVA. Data in this figure correspond to different tissues from some of the animals included in main Fig. 4.

Figure S7. Differences in the loss of type II hair cells (HCII) after exposure to the ototoxic compound, IDPN, as a function of the dose (450 to 600 mg/kg, see legend), zone (central/striola vs periphery) and end-organ (utricle, crista and saccule). The images in the upper-left corner show the use of the Myo7a (red) and calretinin (white) labels to identify HCII. Scale bars = 10 μm. Bar graphs show HCII counts (X±SE) according to end-organ, region within it and IDPN dose. Numbers below the bars indicate numbers of animals. *: p<0.05, significantly different from control group, Duncan's test after significant ANOVA. Data in this figure correspond to different tissues from some of the animals included in main Fig. 4.

Figure S8. Differences in the loss of all hair cells (HC) after exposure to the ototoxic compound, IDPN, as a function of the dose (450 to 600 mg/kg, see legend), zone (central/striola vs periphery) and end-organ (utricle, crista and saccule). The images in the upper-left corner show the use of the Myo7a (red) label to identify HCs. Scale bars = 10 μm. Bar graphs show HC counts (X±SE) according to end-organ, region within it and IDPN dose. Numbers below the bars indicate numbers of animals. *: p<0.05, significantly different from control group, Duncan's test after significant ANOVA. Data in this figure correspond to different tissues from some of the animals included in main Fig. 4.

Figure S9. Relationship between HCI loss and tail-lift angle decrease after ototoxic exposure. Individual data shown here correspond to those shown as group means in main Fig. 2A-C (angles) and Fig. S6 (HCI counts). On each panel, the vertical dashed lines indicate the limits of normal angles (120 degrees, defined by the Control group) and angles in complete absence of vestibular function (70 degrees, defined by the IDPN1000 group). These limit values classified rats into three groups with high, medium and low angles. The p value in each panel indicates statistical significance in HCI counts among these three groups of rats. Double-headed arrows denote significance (p<0.05) of the post-hoc pair-wise comparisons.

Figure S10. Relationship between HCII loss and tail-lift angle decrease after ototoxic exposure. Individual data shown here correspond to those shown as group means in Fig. 2A-C (angles) and Fig. S7 (HCII counts). On each panel, the vertical dashed lines indicate the limits of normal angles (120 degrees, defined by the Control group) and angles in complete absence of vestibular function (70 degrees, defined by the IDPN1000 group). These limit values classified rats into three groups with high, medium and low angles. The p value in each panel indicates statistical significance (p<0.05) in HCII counts among these three groups of rats.

Figure S11. Relationship between all HC loss and tail-lift angle decrease after ototoxic exposure. Individual data shown here correspond to those shown as group means in Fig. 2A-C (angles) and Fig. S8 (all HC counts). On each panel, the vertical dashed lines indicate the limits of normal angles (120 degrees, defined by the Control group) and angles in complete absence of vestibular function (70 degrees, defined by the IDPN1000 group). These limit values classified rats into three groups with high, medium and low angles. The p value in each panel indicates statistical significance (p<0.05) in HC counts among these three groups of rats.

Figure S12. Relationship between HCI loss and air-righting time increase after ototoxic exposure. Individual data shown here correspond to those shown as group means in Fig. 2E-F (times) and Fig. S6 (HCI counts). On each panel, the vertical dashed lines indicate the limits of normal times (170 ms, defined by the Control group) and times in complete absence of vestibular function (350 ms, defined by the IDPN1000 group). These limit values classified rats into three groups with low, medium and high times. The p value in each panel indicates statistical significance (p<0.05) in HCI counts among these three groups of rats.

Figure S13. Relationship between HCII loss and air-righting time increase after ototoxic exposure. Individual data shown here correspond to those shown as group means in Fig. 2E-F (times) and Fig. S7 (HCII counts). On each panel, the vertical dashed lines indicate the limits of normal times (170 ms, defined by the Control group) and times in complete absence of vestibular function (350 ms, defined by the IDPN1000 group). These limit values classified rats into three groups with low, medium and high times. The p value in each panel indicates statistical significance (p<0.05) in HCII counts among these three groups of rats.

Figure S14. Relationship between all HC loss and air-righting time increase after ototoxic exposure. Individual data shown here correspond to those shown as group means in Fig. 2E-F (times) and Fig. S8 (all HC counts). On each panel, the vertical dashed line indicates the limits of normal times (170 ms, defined by the Control group) and times in complete absence of vestibular function (350 ms, defined by the IDPN1000 group). These limit values classified rats into three groups with low, medium and high times. The p value in each panel indicates statistical significance (p<0.05) in HC counts among these three groups of rats.
